# Predicting Peptide HLA-II Presentation Using Immunopeptidomics, Transcriptomics and Deep Multimodal Learning

**DOI:** 10.1101/2022.09.20.508681

**Authors:** Hesham ElAbd, Mareike Wendorff, Tomas Koudelka, Christian Hentschker, Ann-Kristin Kamps, Christoph Prieß, Lars Wienbrandt, Frauke Degenhardt, Tim A. Steiert, Petra Bacher, Piyush Mathur, David Ellinghaus, Uwe Völker, Andreas Tholey, Tobias L. Lenz, Andre Franke

## Abstract

The human leukocyte antigen (HLA) class II proteins present peptides to CD4^+^ T cells through an interaction with T cell receptors (TCRs). Thus, HLA proteins are key players in shaping immunogenicity and immunodominance. Nevertheless, factors governing peptide presentation by HLA-II proteins are still poorly understood. To address this problem, we profiled the blood transcriptome and immunopeptidome of 20 healthy individuals and integrated the profiles with publicly available immunopeptidomics datasets. In depth multi-omics analysis identified expression levels and subcellular locations as import sequence-independent features governing presentation. Levering this knowledge, we developed the Peptide Immune Annotator Multimodal (*PIA-M*) tool, as a novel pan multimodal transformer-based framework that utilises sequence-dependent along with sequence-independent features to model presentation by HLA-II proteins. *PIA-M* illustrated a consistently superior performance relative to existing tools across two independent test datasets (area under the curve: 0.93 vs. 0.84 and 0.95 vs. 0.86), respectively. Besides achieving a higher predictive accuracy, *PIA-M* with its Rust-based pre-processing engine, had significantly shorter runtimes. *PIA-M* is freely available with a permissive licence as a standalone pipeline and as a webserver (https://hybridcomputing.ikmb.uni-kiel.de/pia). In conclusion, *PIA-M* enables a new state-of-the-art accuracy in predicting peptide presentation by HLA-II proteins *in vivo*.

## INTRODUCTION

The classical human leukocyte antigen (HLA) class II proteins are a group of glycoproteins that are mainly expressed on the surface of antigen presenting cells where they present peptides to CD4^+^ T cells. The set of peptides presented by an HLA protein is referred to as the immunopeptidome (1). Polymorphisms in HLA-II coding genes have been genetically linked to a wide range of autoimmune and inflammatory diseases including inflammatory bowel disease (2), multiple sclerosis (3), rheumatoid arthritis (4), and celiac disease (5). Nevertheless, in most cases, potentially causative HLA-peptide interactions remain to be elucidated. Hence, a deeper understanding of factors governing peptide presentation by HLA proteins is of paramount importance to understand disease aetiology and to develop new therapies (1). Furthermore, the ability of HLA-II proteins to present neoantigens to CD4^+^ T cells (6) has made immunopeptidomic profiling crucial for developing novel immunotherapies (7) and cancer vaccines (8).

Given their clinical utility, different methods have been utilized to characterize the set of peptides presented by HLA proteins, *e.g*. competition-based peptide binding assays (9) and peptide microarrays (10). Liquid chromatography-mass spectrometry (LC-MS)-based immunopeptidomics has emerged as the gold-standard method for characterizing the set of peptides presented by HLA proteins *in vivo*, referred to hereafter as the immunopeptidome (11). Despite the widespread popularity of the method, it is still expensive, low-throughput, and time-consuming, thus, computational methods have been developed to predict peptide-HLA interaction *in silico*. Arguably, statistical learning methods such as *NetMHCIIpan* (12), have become the most commonly utilized class of computational methods to predict the binding between peptides and HLA-II proteins. These methods depend on analysing the sequence of presented peptides to learn statistical properties governing presentation by HLA proteins.

The recent rise of deep learning (DL) algorithms and frameworks coupled with the growth of public databases, due to the surge in MS-based immunopeptidomics (11), has fuelled the development of DL methods for modelling peptide-HLA-II interaction. Recently, *Chen et al*. (13) developed *MARIA* using a recurrent neural network based architecture, while *Shao* and colleagues (14) developed *MHCnuggets* employing a similar architecture. Furthermore, *Cheng et al*. (15) developed *BERTMHC* using a Transformer-based architecture while *Graham et al*. (16) developed *BOTA* using a convoluted neural network. Despite the recent breakthroughs which enabled more accurate predictions, different aspects remain uncharacterized, *for example*, the computational efficiency of different methods. For example, *Cheng* and colleagues (15) have recently released *BERTMHC* which is a transformer-based model for predicting peptide-HLA-II interaction. *BERTMHC* is based on *TAPE* (17) which is a large protein sequences model with twelve multi-headed attention layers each with twelve attention heads (15). Hence, predicting the binding between a peptide and an HLA molecule using *BERTMHC*, requires a lot of computational resources and, subsequently, are more expensive in terms of energy consumption and computational infrastructure requirement. Furthermore, the characterization of sequence-independent features that govern presentation by HLA-II proteins *in vivo* remains poorly understood.

To address these limitations, we first generated a large dataset containing the immunopeptidome and the transcriptome of peripheral blood mononuclear cells (**PBMCs**) obtained from 20 healthy donors (**Fig. 1A**). This dataset was combined with publicly available datasets to provide an in-depth analysis of factors governing presentation by HLA-II proteins (**Fig. 1B**). We then used the knowledge extracted from the analysis to develop two novel pan peptide-HLA-II prediction models: (i) peptide immune annotator *(PIA)-S* for sequence-only based predictions (**Fig. 1C**) and (ii) *PIA-M* for multi-modal, *i.e*. multi-omics, based predictions (**Fig. 1E**). To enable a swift integration of multimodal datasets, we developed the <u>om</u>ics <u>li</u>nking toolkit (*OmLiT*) which is a speed-optimized Rust library with a Python binder for pre-processing and preparing multi-omics data that can be used for training or running inferences using the *PIA-M* architecture (**Fig. 1D**). Lastly, we developed a computational pipeline that can be utilized to study peptide-HLA-II interaction from a variety of inputs including complete proteomes and genetic variation data stored in VCF files using the developed models (**Fig. 1F**). Our findings illustrated an important role for sequence-independent features in shaping peptide presentation by HLA-II proteins. Further, our analyses illustrate the utility of deep multi-modal training in improving the predictive accuracy of peptide HLA-II interaction prediction models.

**Figure 1:**
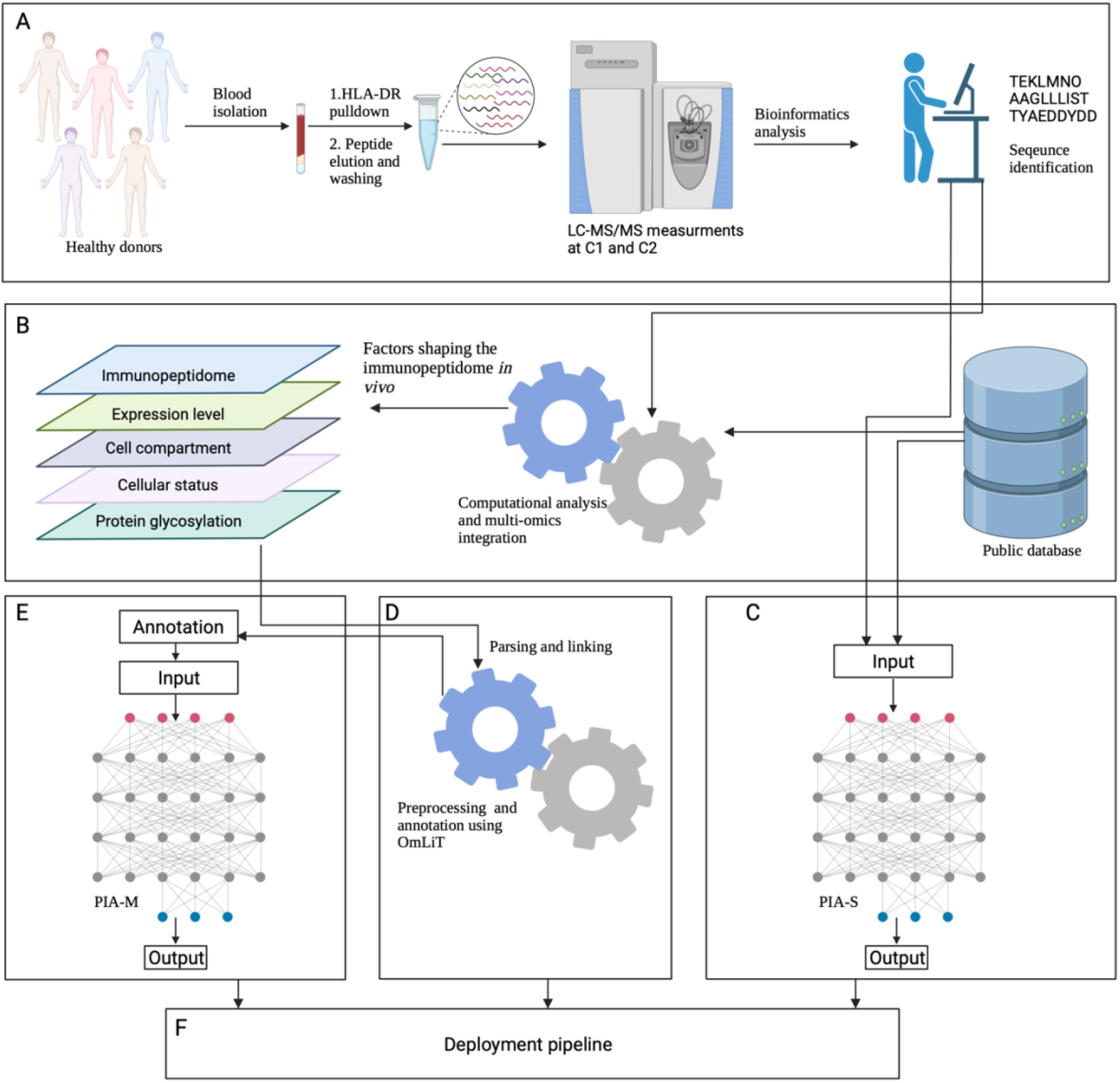
A graphical summary of the study. (**A**) MS-based immunopeptidomics profiling of the study participants. (**B**) Integration with public databases and identifying factors governing peptide presentation in vivo. (**C**) The development of *PIA-S* architecture along with investigating factors governing the predictive performance and the runtime of transformer-based models. (**D**) Developing the omics linking toolkit (*OmLiT*) for preparing and annotating input to *PIA-M*. (**E**) The development of the *PIA-M* model and benchmarking the performance against independent test datasets. (**F**) The development of a pipeline for deploying *PIA-S* and *PIA-M* and enabling large-scale peptide HLA-II interaction characterization. This figure was created by with BioRender.com.

## MATERIAL AND METHODS

### Study design and sample collection

Blood was collected from 27 healthy donors through the University Medical Center Schleswig-Holstein (UKSH) blood bank. All blood donors gave informed consent (Ethics committee *UKSH Kiel*, identifier *D578/18*). For each donor, PBMCs were isolated and quantified using flow cytometry and propidium iodide staining. All 27 donors were used for MS-immunopeptidomic profiling, the blood transcriptome of 25 donors was additionally profiled using RNA-Seq (**Supplementary Methods**). For RNA-Seq, two replicates each containing 1×10^7^cells were isolated per donor. For MS-immunopeptidomic profiling, at least two replicates each containing 1×10^8^cells were utilized. The immunopeptidomes were measured in two proteomics labs, first, at Kiel University referred to as center one (C1) and at Greifswald University referred to as center two (C2). After HLA-DR pulldown, MS-immunopeptidomic profiling and quality-control (at least two replicates per sample), an immunopeptidomic profile for 25 samples that are derived from 23 donors were obtained, where three profiles were measured only at C2, 18 profiles only at C1, and two profiles at C1 and C2 (*i.e*. a total of 4 profiles) (**Table S1)**. After selecting only profiles with at least two replicates, a paired dataset containing the immunopeptidome and the transcriptome of 20 probands (**Table S2)** was obtained.

### Development of *PIA-S* architecture

*PIA-S* is a transformer-based model that is used for predicting peptide-HLA-II interaction using two inputs: the first is the peptide sequence and the second is the HLA-II protein pseudo-sequence (18). Peptides are numerically encoded and are zero-padded into a fixed length array of size 21. Meanwhile, HLA-II pseudo-sequences as defined by *NetMHCIIpan* (18) are extracted and are numerically encoded into a fixed length array of size 34. These two arrays are then concatenated and are fed to a learned embedding layer which projects each amino acid, more specifically, the number corresponding to it, into a learnable encoding space with 32 dimensions. The output of the learned embedding is fed to a stack of self-attention layers each with four attention-heads (19). An average pooling layer is applied along the features axis of the tensor produced by self-attention blocks, reducing the rank of this tensor from three to two. The produced tensor is fed to a feedforward neural network composed of a two multilayer perceptron and a dropout layer (20) to regularize the model and prevent overfitting. The final output is computed using a prediction head with a sigmoid activation function producing a presentation probability.

### Development of *PIA-M* architecture

*PIA-M* is a transformer-based model for predicting peptide-HLA-II interaction using sequence-dependent and sequence-independent features. *PIA-M* receives five inputs: first, the peptide sequence concatenated with the pseudo-sequence of the target HLA-II protein. Second, the expression level of the parent transcript, third, the subcellular compartment of the parent protein, fourth, the context vector which is a vector containing the expression of all coding genes and fifth, the distance to the nearest glycosylation site. *PIA-M* is tightly coupled to the multi-omics linking toolkit library (*OmLiT*) and relies on it for input annotation, pre-processing and encoding. Hence, the input to *OmLiT* is the main entry point to use *PIA-M* either for inference or for training. It is worth mentioning that *OmLiT* and *PIA-M* are integrated into the training pipeline and, more importantly, into the inference pipeline as discussed below. *OmLiT* takes four inputs to prepare a *PIA-M-*compatible input array, first, the source proteome (in most cases this is the human proteome which contains *Uniprot* accessions as keys and protein sequences as values). Second, a table containing HLA-II pseudo-sequences; third, the annotation table which contains detailed annotations of human proteins in different tissues, *e.g*. the expression level of a particular transcript in a particular protein (https://github.com/ikmb/OmLiT). Fourth, a user provided tuple containing peptide sequences, source tissues and HLA alleles. These four inputs are processed by *OmLiT* to produce encoded arrays that can be used for running inference on the model. *PIA-M* depends on two types of blocks to process the inputs: an attention block like *PIA-S* which is used for analysing the input sequence, and a feedforward block for analysing the omics data. The feedforward network is based on a stack of multilayer perceptron (**MLPs**) separated by a dropout layer (20) to reduce overfitting. Scalar values such as the parent expression level and the distance to nearest glycosylation sites are processed using a feedforward layer with only one neuron while high-dimensional inputs, *e.g*. the context-vector which is a vector with ∼16,000 elements, are processed using multiple feedforward layers followed by a dropout layer. The output of the attention-blocks and the feedforward blocks are concatenated and fed to an output prediction head, which produces presentation probabilities.

### Benchmarking the performance of PIA-S and PIA-M using different immunopeptidomic datasets

Three datasets were used for benchmarking the performance of *PIA-S* and *PIA-M*.

1. The first dataset (benchmarking-1) was constructed from the immunopeptidome of three samples that were measured in the current study, with the following HLA-*DR* genotypes: HLA-*DRB1*03:01*/ HLA-*DRB1*11:01*, HLA-*DRB1*11:01*/HLA-*DRB1*11:02* and HLA-*DRB1*07:01*/HLA-*DRB1*09:01* respectively.
2. The second dataset (benchmarking-2) was obtained from *Wang et al*. (21) and contains the immunopeptidome of two HLA-*DR* alleles, (i) *DRB5*01:01* and (ii) *DRB1*15:01*. These two datasets were measured across multiple cell lines, however, for benchmarking *PIA-S* and *PIA-M*, the set of peptides identified from all cell lines was used.
3. The third dataset (benchmarking-3) was assembled from an HLA-DP immunopeptidomic dataset obtained from *Laghmouchi et al*. (22). The dataset contained 13 different immunopeptidomic datasets each of which was obtained from a different HLA-DP protein. To generate peptide sequences, raw MS spectrometry measurements were obtained from PRIDE (23) under the accession PXD030591 and were subsequently processed using *MHCquant* (24) (**Supplementary Methods**)

Performance metrics, namely, the area under the receiver operator curve (ROC AUC) and the area under the precision recall curve (PR AUC) were calculated using *TensorFlow* (25) and *Scikit* library (26).

## RESULTS

### Sequence-independent features shape HLA-II immunopeptidomes

We started by characterizing the HLA-DR immunopeptidome of different healthy individuals using blood samples (**Material and Methods**). We focused on HLA-*DR* as it has been previously shown to be the dominantly expressed HLA-II gene (27) and on blood as it provides a relatively accessible tissue to obtain biologically-resemblant samples as opposed to cell lines. Using two biological replicates per sample (each utilising 1×10^8^ PBMCs), we identified on average 3,265 unique peptides and 970 unique proteins per sample (**Fig. 2A**). The identified peptides had the expected length distribution for HLA-II peptides with most peptides being 13- to 17-mers (**Fig. S1**).

**Figure 2:**
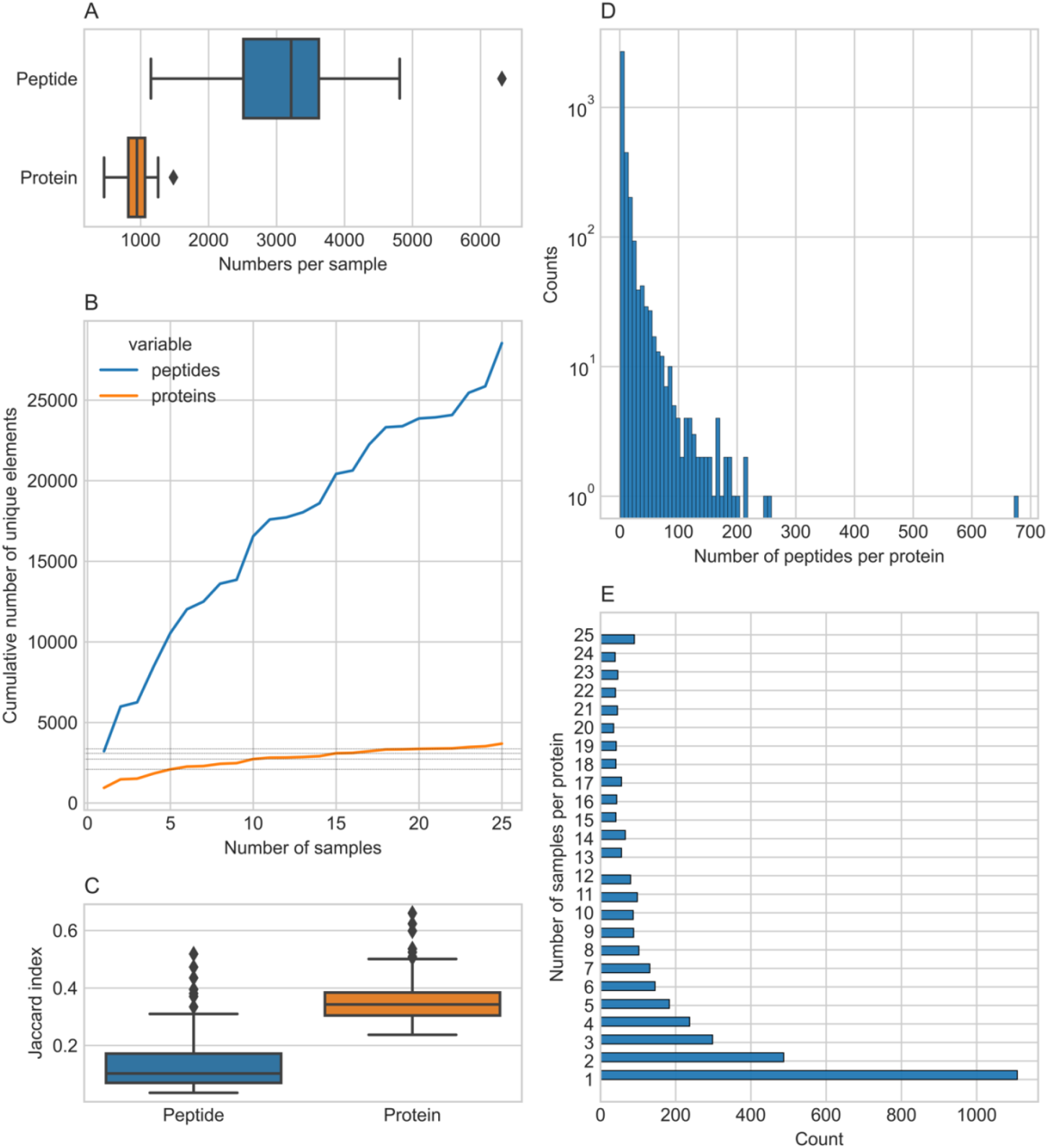
An overview of the data derived from the samples characterized in the current study. As mentioned in the **Material and Methods**, the blood HLA-DR immunopeptidome of 23 healthy probands were profiled, however, two samples were independently measured at centre 1 and centre 2, hence the results here summarise findings of 25 samples obtained from 23 probands. (**A**) The distribution of unique peptides and proteins per sample. (**B**) The cumulative number of peptides and proteins across all samples where grey dash-lines highlight the number of unique proteins after 5, 10, 15, and 20 samples, effectively, illustrating the plateauing in the number of unique proteins with an increased number of samples. (**C**) The distribution of Jaccard indices between samples computed pairwise at the peptide-level and the protein level. (**D**) A histogram of the number of peptides-per-protein. (**E**) The frequency of protein presentation among different samples, *i.e*. the number of samples where a particular protein was observed or presented by an HLA-DR molecule.

Next, we looked at the cumulative number of peptides and proteins among samples. As seen in **Fig. 2B**, the number of unique peptides continuously increases with each sample following a, more-or-less, logarithmic growth where the number of new proteins identified plateaus after ∼15 samples. This suggests that different samples present different peptides, nevertheless these peptides originate from a subset of the human proteins. In essence, this implies that different HLA-DR proteins present different peptides, however, these peptides originate from a smaller set of enriched proteins.

To characterize this further, we computed the Jaccard index (defined as intersection divided by the union of peptides and/or proteins identified between two samples) among different independently collected samples at the peptide and the protein level. As seen in **Fig. 2C**, the samples, are more similar, *i.e*. have a higher Jaccard index (J-index), at the protein level than at the peptide level confirming previous observations. Nevertheless, it worth mentioning that comparing the J-index between peptides and proteins is challenging because the difference in the number of unique peptides and protein per sample. Nonetheless, the computed scores are spread over a long range of values at the peptide level (J-index ∈ [0.0356,0.5186]) and at the protein level (J-index ∈ [0.2374,0.6603]). To get a better idea of factors governing similarity of presentation between samples, we compared the HLA-*DR* genotype of sample-pairs exhibiting a high score at the peptide-level (J-index> 0.3) and at the protein-level (J-index > 0.5) (**Table S3**). As seen in the table, all similar samples share, at least, one HLA-*DR* allele. To measure the impact of shared HLA-*DR* background on the J-index, we calculated the correlation between the J-index and the number of shared alleles (two-fields resolution) at the peptide-level (**Fig. S2A**) and at the protein-level (**Fig. S2B**) across all unique sample-pairs. As seen in the figure, there is a positive and significant relationship between the number of shared HLA-II alleles and similarity at the immunopeptidome level. Furthermore, at the protein level, samples with completely different HLA-*DR* alleles, exhibit a relatively high level of overlap (J-index∼0.3). This suggests that next to the HLA proteins binding preferences and peptide sequences, additional features, play a role in shaping the landscape of HLA-II immunopeptidomes.

To study this further, we looked at the number of peptides per protein among all samples. As seen in **Fig. 2D**, most proteins are identified by only one peptide. Meanwhile, a small proportion of proteins are supported by tens to hundreds of peptides. Furthermore, by looking at the frequency of protein presentation among samples (**Fig. 2E**), we observed that ∼30% of proteins are identified once, *i.e*. in only one sample, while the remaining ∼70% are observed among different samples with different frequencies. These findings suggest that other factors beyond peptide sequence contribute to shaping the HLA-DR immunopeptidomic landscape. To investigate these factors, we compared the average expression of the presented proteins to non-presented, *i.e*. non-observable proteins (**Supplementary Methods**). As seen in (**Fig. 3A**) presented proteins have, on average, a higher expression level than non-presented (non-observed) proteins, confirming previous findings by *Chen et al*. (13). Furthermore, we identified a positive correlation between the frequency of protein presentation among samples and the parent transcript expression level (**Fig. 3B**). Next, we investigated the impact of the cellular compartment on shaping the immunopeptidomic landscape. By focusing on the proteins identified in all samples, we identified an enrichment in cellular compartments related to vesicles trafficking, plasma membrane, secreted proteins and, HLA-II proteins loading machinery using gene ontology enrichment analysis (**Fig. 3C, Fig. S3**). As seen in **Fig. 3B**, proteins identified in all samples exhibit, on average, higher expression levels relative to proteins identified in only one sample. Nevertheless, the expression level of these genes showed a wide range of variation, suggesting that abundance and compartment are also contributing to peptide presentation. These findings suggest an important role for sequence-independent protein features, *e.g*. expression level which is a proxy for abundance, along with the cellular compartment, in modulating the set of self-proteins presented by HLA-II proteins *in vivo*.

**Figure 3:**
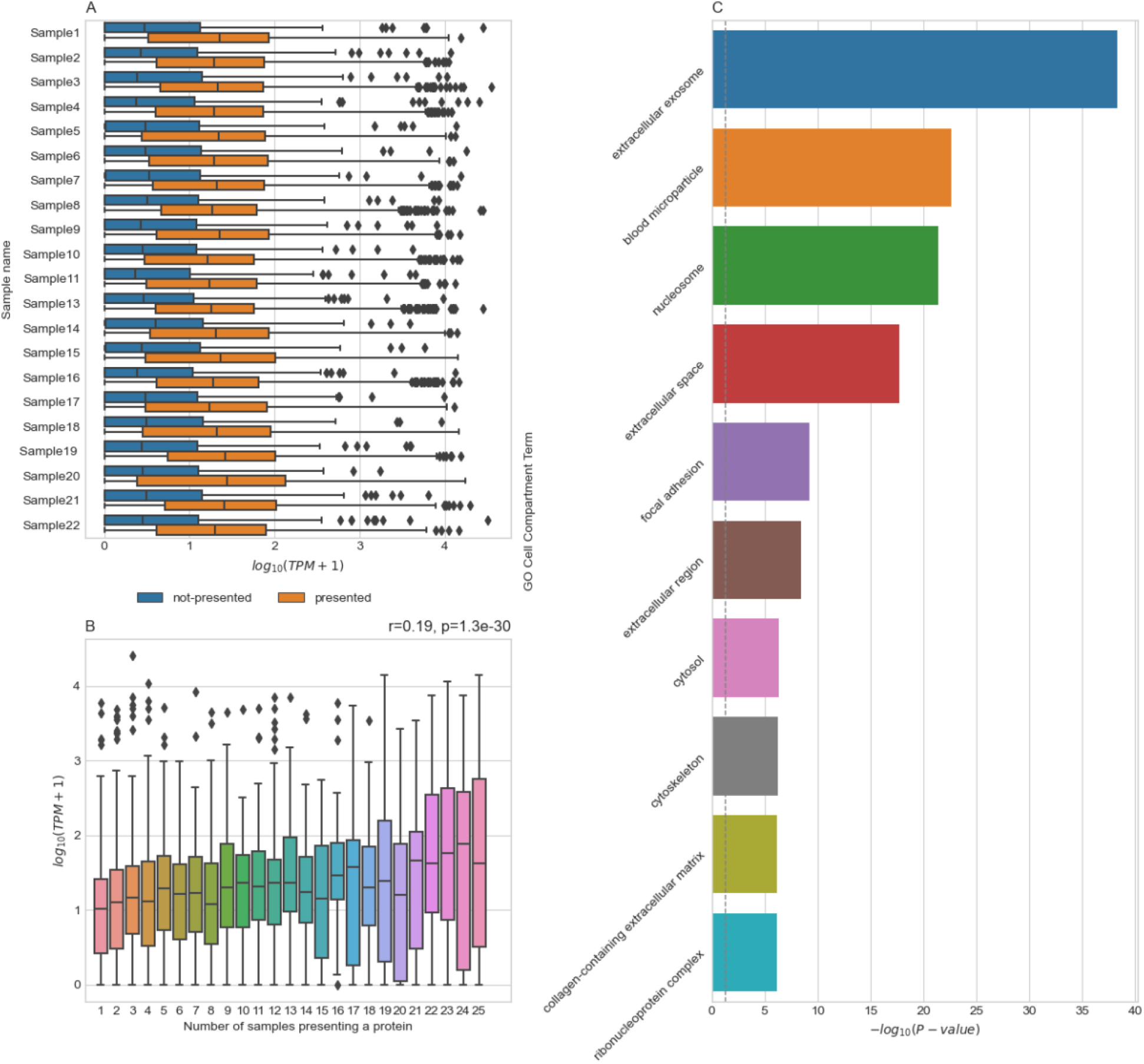
An overview of sequence-independent features governing peptide presentation by HLA-II proteins. (**A**) A comparison between the expression levels (in transcripts per million (TPM)) of presented proteins and proteins that were not observed. Not-presented proteins were generated for each sample by sampling from the set of not-observed proteins a subset of genes with the same cardinality, *i.e*. the number of genes, as the set of positive proteins. It is worth mentioning that Sample 14 and Sample 15 share the same transcriptomic profile as they were obtained from the same donor and measured at two different centres, further the expression data was obtained using the paired dataset (**Material and Methods**) hence, only 20 samples are only shown. (**B**) The correlation between the number of samples presenting at least one peptide from a particular protein and the protein’s transcript expression level as defined in the Human Protein Atlas database (33). The Pearson correlation coefficient was computed between the two variables identifying a significant (p=1.3×10^−30^) but moderately positive relationship (r=0.19). (**C**) A gene ontology enrichment analysis (GOEA) for the 90 proteins presented in all samples focusing on the top 10 subcellular compartments. Here, the dashed grey line represents the significance threshold of (-*log*_10_(0.05) = 1.3 after false discovery rate (FDR) correction as implemented in GOATOOLS. GOEA was conducted using GOATOOLS version 1.1.6 (34) and IPTK (35).

To validate these findings, we used a large immunopeptidomics dataset that was recently described by *Reynisson et al*. (28). First, we tried to validate the pattern observed in **Fig. 2E** by analysing the number of peptides per protein across all the studies included in the public database. As seen in **Fig. S4A**, most proteins are supported by one or few peptides with the minority supported by hundreds to thousands of peptides. Indeed, proteins with most peptides in our own dataset **Fig. S5** are also highly presented, *i.e*., are supported by many peptides, in the public dataset (**Fig. S6**). Next, we tested the hypothesis that protein length might correlate with the number of peptides per protein (**Fig. S4B**). As seen in the figure, there was no significant correlation between protein length and peptide presentation (p=0.2). Then, we aimed at replicating the pattern observed in **Fig. 2B** by analysing the cumulative number of peptides and proteins among different studies (**Fig. S4C**). A pattern resembling **Fig. 2B**, can be observed where the number of peptides keeps growing by increasing the number of experiments meanwhile the number of proteins plateaus, or at least grow logarithmically, after a few experiments. This suggests that besides the sequence length other sequence-independent features such as protein abundance and availability to the HLA-II loading machinery are not only pivotal in shaping protein presentability, *i.e*. at least one peptide from the protein is presented, but also the density of presentation, *i.e*. the number of peptides per protein.

### Balancing the predictive performance and the computational efficiency of different transformer-based models

So far, we have focused on identifying factors governing peptide presentation *in vivo*. However, as explained above, it is experimentally infeasible to characterize the interaction between HLA-II proteins and the vast amount of potential peptide candidates. Hence, *in silico* peptide predictions models are needed for modelling peptide-HLA-II interaction. Given, the superior performance shown by *BERTMHC* (15), we focused only on transformer-based models.

We started by quantifying the impact of the model complexity, specifically, the depth (number of layers) and the number of attention heads per layer, on the inference time of a transformer-based model. To do this, we created different models following *PIA-S* architecture (**Fig. 5A**; **Material and Methods**) with different number of layers and different number of attention heads per layer. We measured the inference runtime using a mock dataset representing one million peptide-HLA-II pairs. As seen in **Fig. 4A**, increasing the number of layers and the number of heads had a strong impact on the runtime, *e.g*. the inference time of a one-layer model with one attention head is ∼5 seconds meanwhile the runtime of a twelve-layers model with 16 attention-heads is ∼111 seconds.

**Figure 4:**
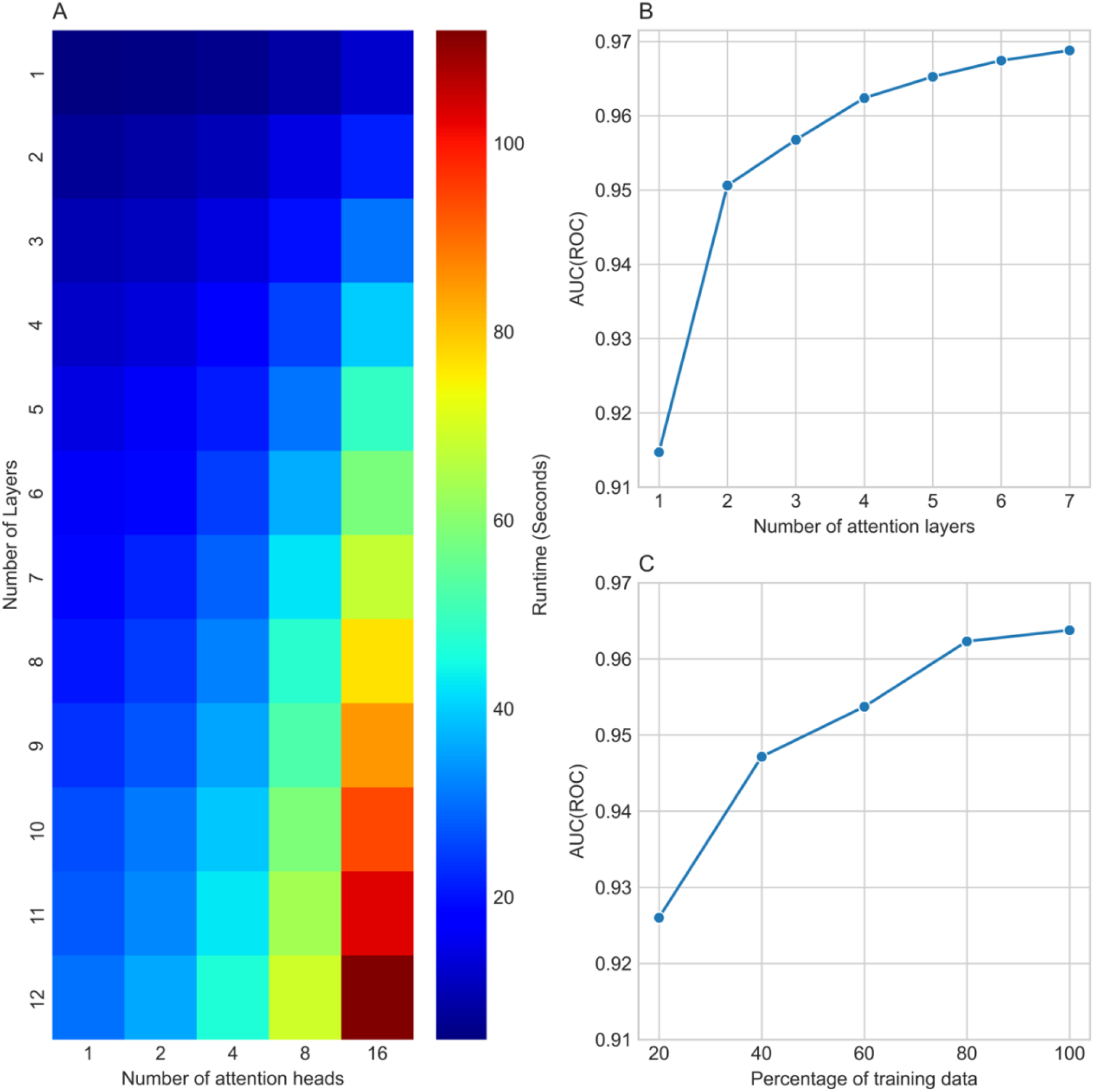
Factors shaping the predictive performance and the inference runtime of transformer-based models. (**A**) A heatmap showing the impact of the number of self-attention layers and the number of attention heads per layer on the runtime of the models where the runtime was averaged across ten different replicates. (**B**) The impact of number of attention layers on the predictive accuracy of transformer-based models (with 4 attention-heads) on the validation dataset. (**C**) The impact of the amount of training data on the model performance on the validation dataset. Here, a model with three multi-attention layers each with four attention heads was used for training the models.

**Figure 5:**
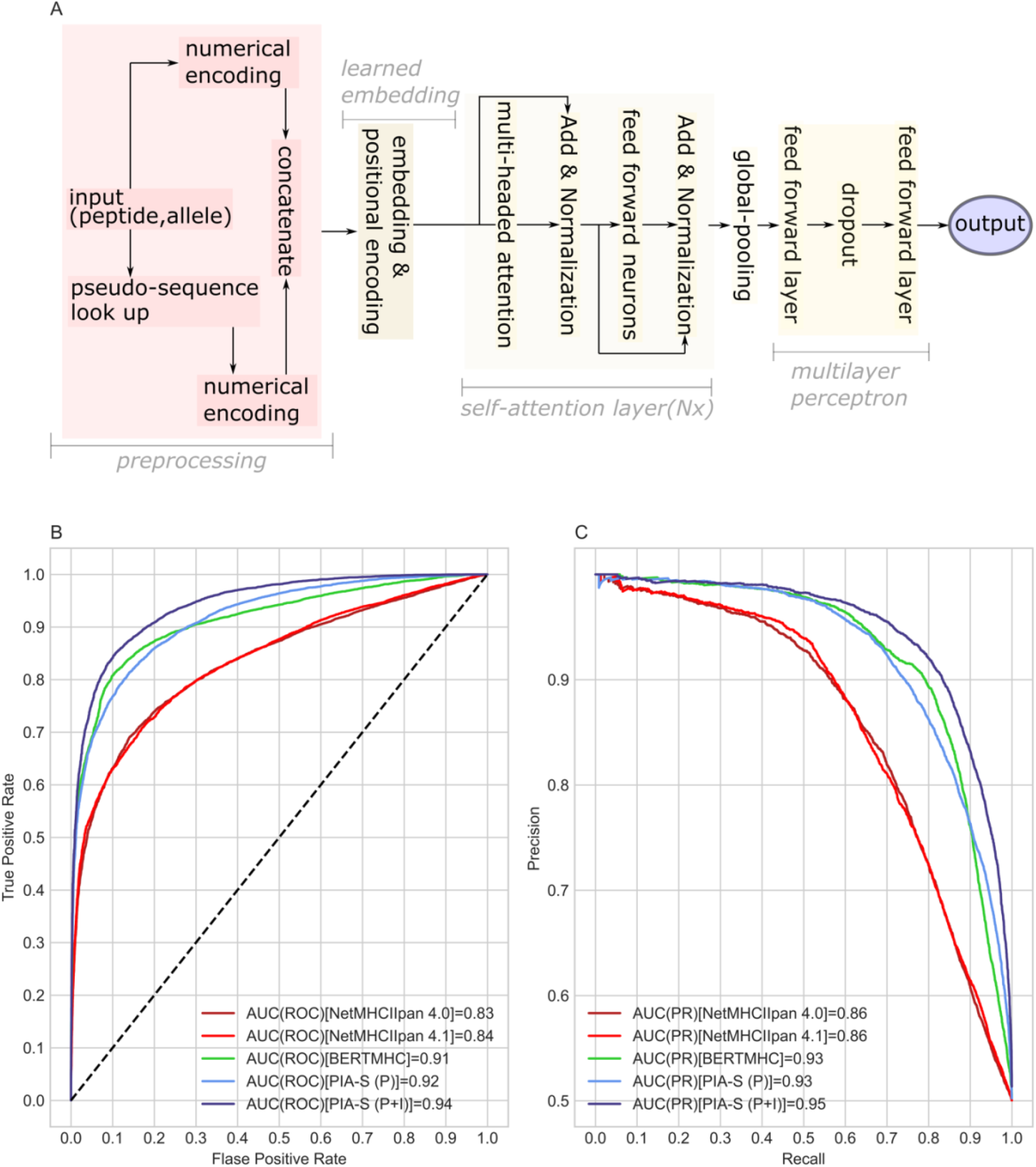
The performance of *PIA-S* against different publicly available prediction methods. (**A**) The blueprint of *PIA-S* (**Material and Methods**) where, briefly, input peptides and pseudo-sequences are tokenized and then concatenated together. Next, the resulting sequence is fed to a learned embedding layer and subsequently to a stack of self-attention layers. Lastly, the resulting tensor in pooled through a global-average pooling and is fed to a multilayer perceptron to calculate the presentability score. The performance of *PIA-S* was benchmarked against *NetMHCIIpan 4.0* (12), *NetMHCIIpan 4.1* and *BERTMHC* (15) using three HLA-DR immunopeptidomics test datasets that were obtained in the current study. (**B**) Shows the receiver operating curve (ROC) along with the area under the curve (AUC(ROC)) using the predictions on the three datasets combined. (**C**) Shows the precision-recall (PR) curve along with the area under the curve (AUC(PR)) using the predictions on the three datasets combined.

**Figure 5:**
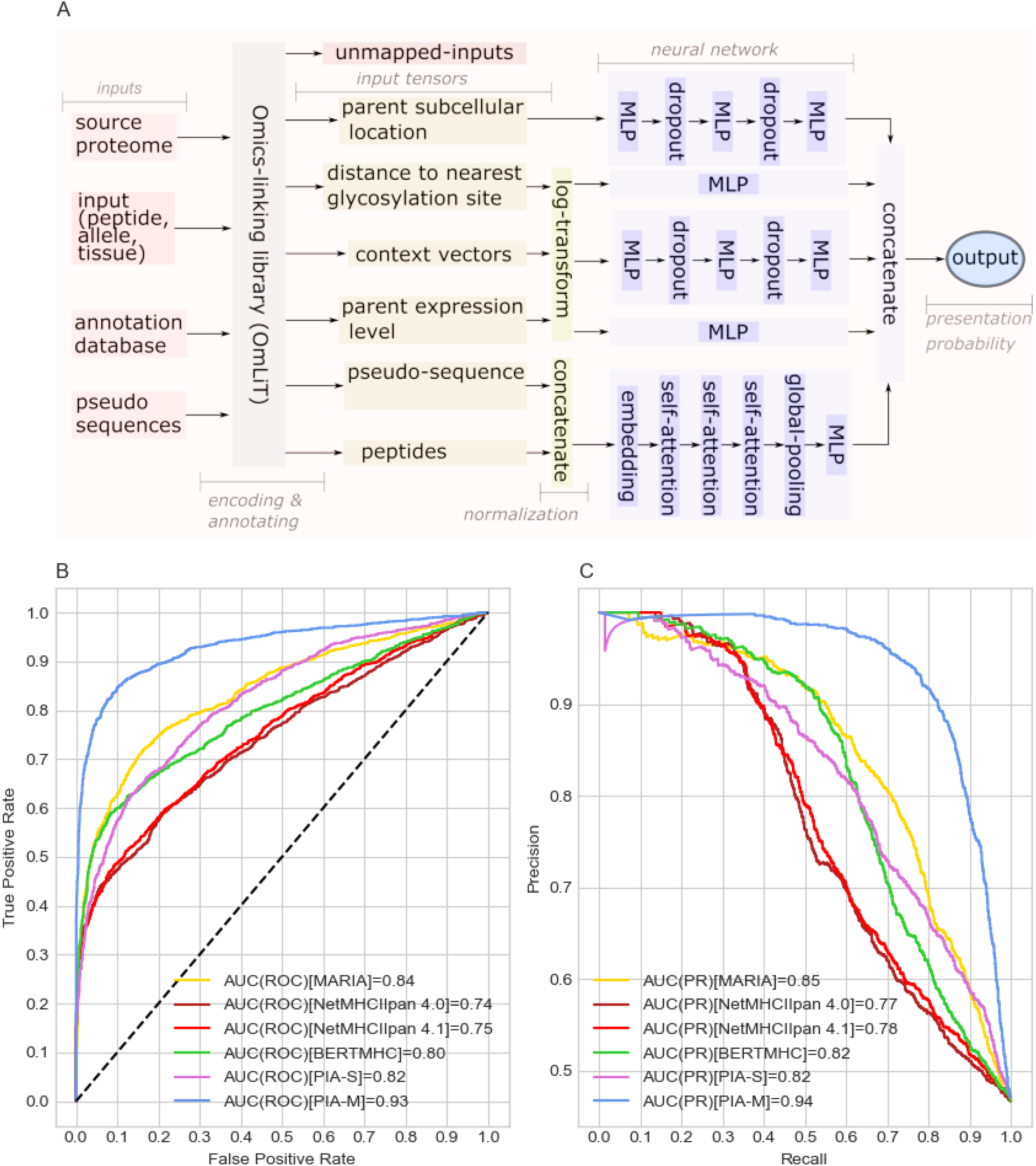
The performance of PIA-S and PIA-M on an independent test dataset containing the immunopeptidome of two HLA-DR alleles, namely: HLA-*DRB5*01:01* and HLA-*DRB1*15:01* obtained from *Wang et al*. (21). (**A**) A description of the *PIA-M* architecture (**Material and Methods**) illustrating the major components, namely the *OmLiT* library which is used for preparing and annotating the input and the neural architecture of *PIA-M* where different omics layers are modelled and combined to predict peptide presentation by HLA-II proteins. (**B**) The receiver operating curve (ROC) of the predictions generated by different tools on the combined dataset of the two alleles, meanwhile (**C**) shows the precision-recall curve (PR) curve.

Next, we tried to quantify the impact of the model’s size on its predictive performance, *i.e*. the accuracy of predicting peptide HLA-II interaction. To this end, we used the public dataset described above to train different *PIA-S* based models with an increasing number of layers (**Material and Methods**). As seen in **Fig. 4B**, a model with one layer was already able to achieve an impressive performance with AUC ROC close to ∼0.91. Increasing the number of layers also increased the predictive performance, which then plateaued at approximately five layers. After that, increasing the number of layers had only a marginal impact on improving the performance. This was independent of the number of training epochs (**Fig. S7**). Subsequently we investigated the effect of increasing the training data on improving the model performance (**Fig. 4C**). Increasing the amount of training data improved the model performance, as expected.

### *PIA-S* achieves state-of-the-art performance on sequence only peptide HLA-II interaction prediction

We trained a *PIA-S* architecture model with only three multi-headed attention layers each with four attention-heads using two datasets, first, the publicly available dataset by *Reynisson et al*. (28) (**Supplementary Methods**). Second, using *Reynisson et al*. (28) dataset and the HLA-DR immunopeptidome of 22 samples (obtained from 20 individuals) measured in the current study. Then, we benchmarked the performance of these models against an independent test dataset composed of the HLA-DR immunopeptidome of three samples measured in the current study (benchmarking data set 1, **Material and Methods, Table S4**). As seen in (**Fig.5B & Fig.5C**), *PIA-S* with three self-attention layers achieved a comparable performance to *BERTMHC* (15) which has 12 attention layers confirming previous findings where increasing the number of layers mainly increases the inference runtime while providing infinitesimal improvement in accuracy.

Interestingly, transformer-based models, *i.e. BERTMHC* (15) and *PIA-S* illustrated a superior performance to *NetMHCIIpan* (12). Here, two versions of *NetMHCIIpan* were used, namely 4.0 and 4.1, the former has been recently described by Reynisson *et al*. (12) and is trained using the same dataset used for developing *BERTMHC* and *PIA-S (P)*. Meanwhile, the latter is an unpublished version that was trained on extended datasets. As seen in (**Fig.5B & Fig.5C**), *NetMHCIIpan 4.1* achieved a slightly higher performance relative to *NetMHCIIpan* 4.0 illustrating a marginal improvement in performance with increasing the data. However, the performance of the two versions was inferior to transformer-based models, arguably showing the need for a complex neural network architecture to capture the complexity of large-scale immunopeptidomics datasets.

### Deep Multi Modal training enables accurate and robust identification of peptide presentation by HLA-II proteins

As discussed above, different sequence-independent factors govern peptide presentation by HLA-II proteins *in vivo*, for example, the expression level of parent transcripts and the subcellular location of proteins. However, most publicly available predictions tools except *MARIA* (13) predict peptide-HLA-II interaction based only on peptide sequences and HLA protein pseudo-sequences. Hence, these tools are not modelling the impact of sequence-independent features on peptide presentation which might hinder the performance of these models. Also, *MARIA* (13) utilizes the expression level and proteolytic digestion along with sequence-dependent features to model peptide-HLA-II interaction.

To characterize the impact of modelling different sequence-independent features relative to sequence only models, we trained different prediction models using data derived from two homozygote samples, namely, HLA-*DRB1*04:01* and HLA-*DRB1*15:01* (**Material and Methods**). As seen in **Fig. S8**, training the model with the four sequence-independent features: *i.e*. (exp +SC+Context+D2G) greatly, improve the performance relative to the baseline of modelling sequence-only features.

Next, we utilized the results to develop the peptide immune annotator multimodal (*PIA-M*) (**Fig. 6A**) and trained it using the same dataset used for training *PIA-S* as described above (**Supplementary Methods**). However, a major hurdle was the efficient annotation and encoding of millions of training examples. To address the problem, we developed the omics linking toolkit (*OmLiT*) library (**Supplementary Methods; Fig. S9**). To benchmark the performance of *PIA-M* against publicly available tools, we utilized two independent publicly available datasets that were recently published and hence were not used for training neither of the models (**Material and Methods, Benchmark dataset 2-3, Table S5 and S6**). As seen in (**Fig. 6B and Fig. 6C; Benchmark dataset 2**), *PIA-M* demonstrated superior performance to all sequence-only models and to MARIA (13). Interestingly, the latter (13) is also a multimodal model that achieved a superior performance to all sequence-only models including *PIA-S*. Thus, demonstrating the importance of modelling sequence-independent features to improve the modelling accuracy of peptide HLA-II interaction. Furthermore, we benchmarked the performance of *PIA-M* against sequence-only models using **Benchmark dataset 3** which contain the immunopeptidome of 13 different HLA-DP proteins (**Fig. 7)**. As seen in the figure, *PIA-M* illustrated a consistent superior performance across the immunopeptidome of the 13 samples (**Fig. 7A**) and on the combined dataset (**Fig. 7B & Fig. 7C**). Thus, confirming the superior performance of *PIA-M* and illustrating the advantage of integrating sequence-independent features. Interestingly, *PIA-S* has demonstrated a comparable performance to *NetMHCIIpan 4.1* and *NetMHCIIpan 4.0* on this test dataset, suggesting that the superior performance of transformer-based models might be specific to HLA-DR only. This is possibly a result of the bias in the training datasets that is mostly derived from HLA-DR, nevertheless, the performance of *PIA-S* is still comparable to *NetMHCIIpan 4.1* on HLA-DP and superior on HLA-DR proteins.

**Figure 6:**
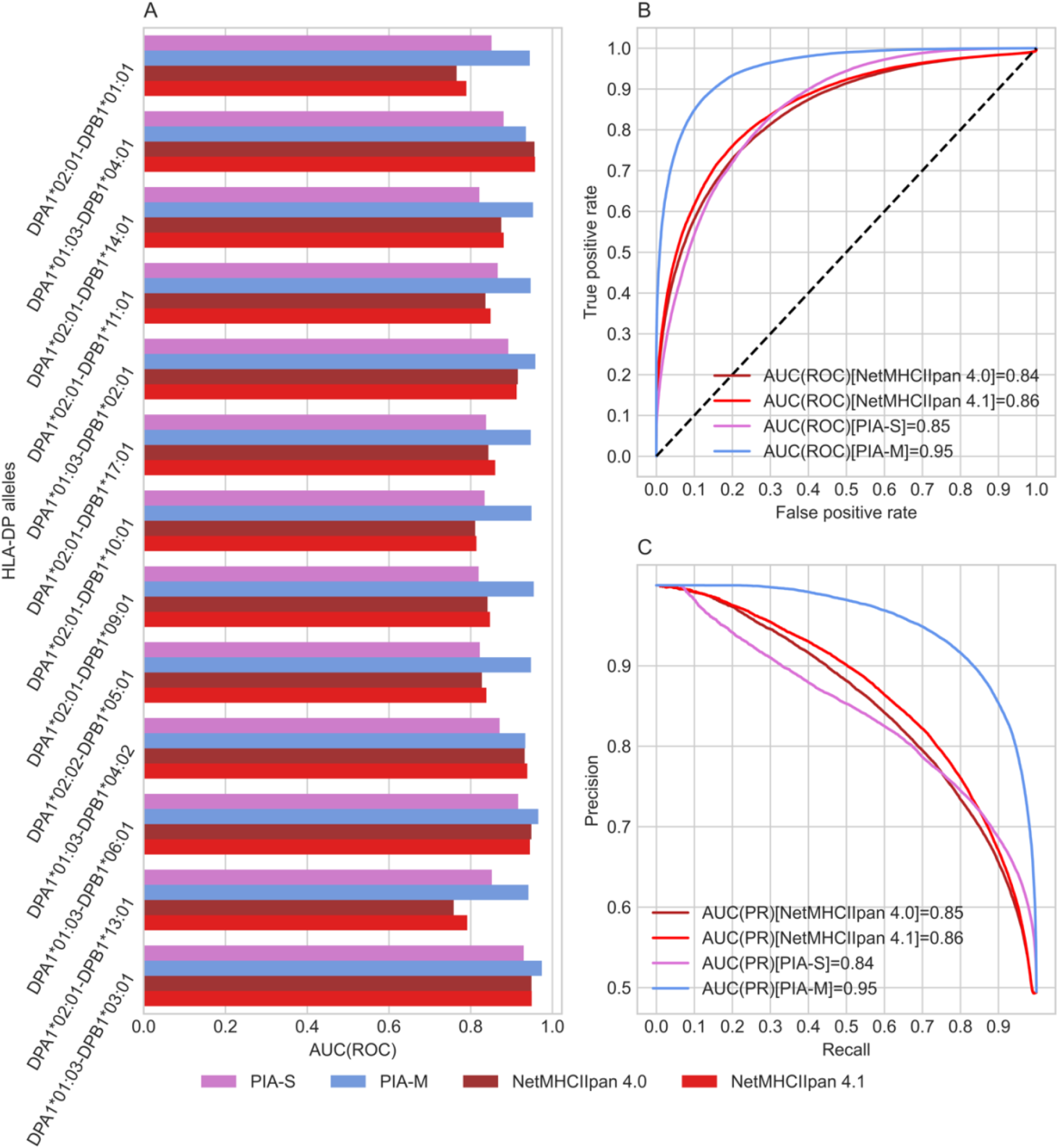
Benchmarking the performance of *PIA-S* and *PIA-M* against *NetMHCIIpan 4.0* (12) and *NetMHCIIpan 4.1* using peptide-elution datasets covering different HLA-DP alleles. The raw files were obtained from PRIDE (23) under the accession *PXD030591* as provided by *Laghmouchi et al*. (22), subsequently, peptides were identified from each allele using *MHCQuant* (24) (**Material and Methods**). (**A**) The predictive accuracy of the four tools on elution data from proteins encoded by HLA-DP proteins (y-axis) where accuracy was measured using the area under the receiver operating curve AUC (ROC) (x-axis). (**B**) The receiver operating curve (ROC) of the predictions of the four tools on the combined dataset of all HLA-DP alleles meanwhile, (**C**) shows precision-recall curve (PR-curve).

**Figure 7:**
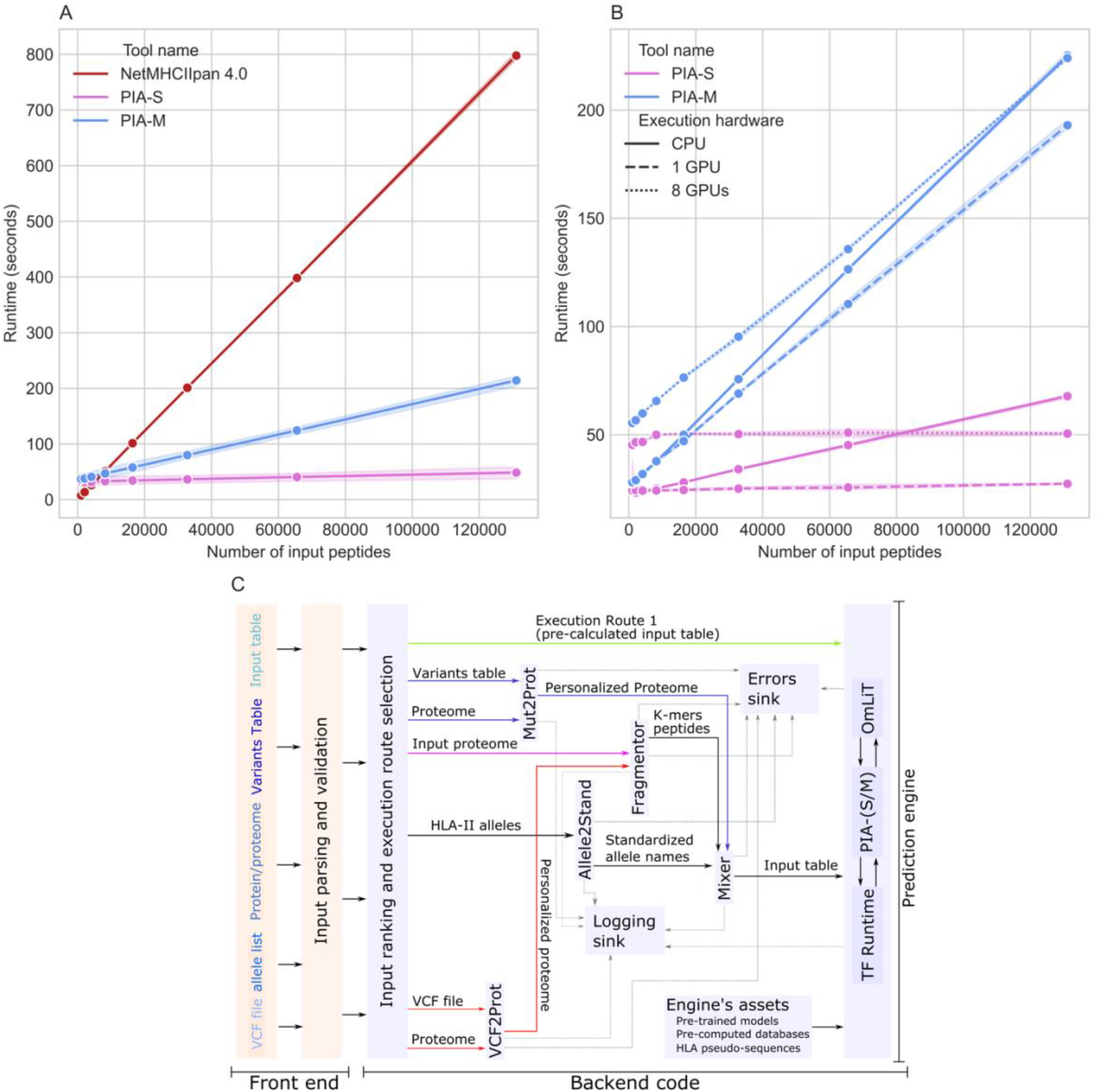
Comparing the execution speed of *NetMHCIIpan 4.0, PIA-S* and *PIA-M*. (**A**) A comparison between the run time of the three tools with an increasing number of input peptides averaged across different execution hardware (**B**) The run time of *PIA-S* and *PIA-M* using different hardware, namely, a CPU, a single GPU and eight GPUs with different inputs. In both A and B, the runtime of each input was measured across three independent replicates. (**C**) A schematic diagram of the developed webserver and the *PIA-P* pipeline focusing on the backend code used for parsing different inputs and scaling-up the execution of peptide-HLA-II interaction. The front-end focuses on parsing user inputs, *e.g*. validating input formats and the correct combination of inputs as discussed below. The pipeline expects four input categories: first using ‘input table’ which is a table that contains the peptide sequence along with the HLA alleles and optionally the tissue name. As the peptide sequence have been generated this table is fed directly to the prediction engine to estimate the interaction between peptides and HLA-II proteins. Second, using a FASTA file containing protein sequences, *i.e*. proteomes, and a list of alleles, here, the expected aim is to characterise the interaction between each possible k-mers peptide in the FASTA file and the protein product of each allele in the provided list of alleles. Here, the input proteome is first fragmented using the *Fragmentor* into fixed-length peptides through a sliding window approach. Next, the names in the list of alleles are parsed using *Allele2Stand* and are combined with the generated list of peptides to generate an input table that is subsequently fed to the prediction engine. The third and the fourth categories are conceptually similar to each other where the aim is to first generate sample-specific proteomes using the reference proteome and sample specific genetic data. Genetic data are supported in two formats, first, variant calling format (VCF) and tab-separated files (TSV). In case of the former, sample specific proteome sequences are generated using *VCF2Prot* (36) meanwhile, in the latter, the *Mut2Prot* is used to do this task. Once sample-specific proteins sequences have been generated they are fragmented using the *Fragmentor* and the generated list of peptides is processed as described above. Lastly, the prediction engine is composited of three main parts, first, the *OmLiT* library which is used for parsing and encoding the input. Second the trained models, namely, *PIA-S* and *PIA-M* which are used for estimating the interaction between peptides and HLA-II proteins. Lastly, the TensorFlow runtime which is used for managing the execution of *PIA-S*/*PIA-M* on the available computational infrastructure, *e.g*. using GPUs.

### *PIA-S* and *PIA-M* have a superior execution speed relative to *NetMHCIIpan*

Motivated by the superior performance of *PIA-M* on multiple independent test datasets we aimed at benchmarking the execution speed of *PIA-S* and *PIA-M* against *NetMHCIIpan 4.0* as it is one of the commonly utilized tools (**Supplementary methods**). As seen in **Fig. 8A**, *PIA-S* and *PIA-M* achieved a significantly faster execution speed relative to *NetMHCIIpan*, indeed the difference in the execution speed widen with larger input size, illustrating the utility of *PIA-S* and *PIA-M* in large-scale screening applications. For example, with 131,072 peptides (2^17^) *NetMHCIIpan 4.0* utilized ∼800 seconds to predict the interaction against a single HLA-II protein. Meanwhile, for the same input *PIA-M* utilized ∼220 seconds, *i.e*. 4 times faster, and *PIA-S* utilized ∼70 seconds, *i.e*. ∼10 times faster. Thus, illustrating not only a superior predictive performance but also a faster execution.

Given that *PIA-S* and *PIA-M* are implemented in TensorFlow (25), they can run on different hardware, for example, CPUs and GPUs, hence, we benchmarked their execution time on different systems (**Fig. 8B**). As seen in the figure, for an average number of peptides (∼100,000-1000,000) a single GPU greatly decreases the runtime, meanwhile, using multiple GPUs is associated with a slower execution time, given the overhead associated with copying and replicating the data and the models across multiple GPUs. Meanwhile, utilizing a CPU-only achieved a slower but practical runtime where the interaction between 131,072 peptides and an HLA-II proteins was predicted in ∼1 minutes and ∼4 minutes by P*IA-S* and *PIA-M*, respectively.

Lastly, to enable an automatic and a swift utilization of the trained models, *i.e. PIA-M*/*PIA-S*, for predicting peptide HLA-II interaction, we developed the peptide immune-annotation pipeline (*PIA-P*) (**Fig. 8C**). The pipeline is currently implemented as a collection of Python scripts that are connected using a Bash script, thus, enabling the pipeline to be utilized on any Linux-based machine with or without GPUs. Conceptually, the pipeline is assembled from three logical components. First, a pre-processing component, which processes different user input formats and generate a list of input peptides, *e.g*. using proteomes stored in a FASTA file and genetic variation data stored in *VCF* files or simple text table. Second, mixing where the names of HLA alleles are parsed and are combined with the generated list of peptides to provide a unified internal representation that is later used by the third component, the prediction engine. The latter is used to calculate the interaction between each input peptide HLA-II pairs using *PIA-S* or to annotate and encode the input with *OmLiT* and run predictions using *PIA-M*.

As stated above, the developed model can run swiftly without a GPU, however, with large input-size, using GPUs greatly improve the execution speed. Hence, to increase the accessibility of the models we developed a web server that runs the developed pipeline, *i.e. PIA-P*, using GPU-accelerated computational infrastructure and allow users to freely utilized all the features of the pipeline using a user-friendly web interface (**Fig. S10**).

## DISCUSSION

Here, we tried to disentangle factors governing presentation by HLA-II proteins, and subsequently, model these factors using deep learning. Beyond the sequence-binding preferences of each HLA alleles, our analysis highlighted some sequence-independent features that contribute to presentation by HLA-II proteins. One of these factors is the transcriptomic landscape of the cell, where highly expressed genes are more likely to be presented by HLA-II proteins than lowly expressed genes which confirms previous findings by *Chen et al*. (13). We also found that proteins derived from highly expressed genes are more likely to be presented by different individuals than proteins derived from lowly expressed genes. However, these results shall be interpreted with caution as we only have a partial picture of the presented immunopeptidome. Indeed, our ability to detect the immunopeptidome is upper bounded by the detection limit of the mass spectrometry. Hence, we cannot reject the hypothesis that lowly expressed genes might be presented at a lower level by HLA proteins *in vivo* which could be detected by more sensitive instruments. However, current data supports a positive relationship between the expression level and presentability by HLA-II proteins.

Interestingly, we also detected an enrichment in extracellular, secreted and vesicle associated proteins in HLA-II immunopeptidomes which supports the current model for HLA-II proteins as presenters mainly for extracellular proteins (29). Despite the importance of sequence-independent features in shaping peptide presentation *in vivo*, most publicly available tools for predicting peptide HLA-II interaction, *e.g. NetMHCIIpan* (12), *BERTMHC* (15) or *MHCnuggets* (14) are based solely on the peptide sequence which might limit the predictive performance of these methods. To address this problem, we developed *PIA-M* which predicts the presentation given sequence-dependent features, *i.e*. the peptide sequence, along with sequence independent features, *e.g*. the transcriptomic landscape, subcellular location and the glycosylation status of the protein. *PIA-M* achieved state-of-the-art performance over its sequence-only version, *PIA-S*, and publicly available sequence-only models.

Annotating all input peptide sequences with sequence-independent features, such as expression or subcellular location is challenging from a biological and a computational perspective. Starting with the former, the majority of immunopeptidomics experiments has focused mainly on self (29) and neoepitopes (30) presentation with few notable exceptions that focused on microbial presentation by HLA-II proteins, *e.g*. Graham *et al*. (16) studying presentation of micro-organisms in mice-derived B cell lines. Thus, our understanding of sequence-independent features governing presentation by HLA proteins is biased toward self-peptide presentation. Hence, a more experimental characterization of microbial presentation by HLA-II proteins would be needed to discover and learn sequence-independent features governing microbial and foreign peptide presentation by HLA-II proteins *in vivo*.

Despite the superior performance of multimodal models, they are more computationally demanding relative to pure sequence models not only due to the complexity of the model but also due to the complex pre-processing and annotation needed. Here, we mitigated this problem by first, offloading the computationally heavy tasks to a speed-optimized code written in Rust, *i.e. OmLiT* library. Second, through generating a database covering only human proteins. Beyond improving computational efficiency, the annotation database has been also restricted to humans as high-quality multi-omics data could be only obtained for human proteins as discussed above.

Immunopeptidomics datasets contain only positive peptides, *i.e*. the binders, hence, negative examples have to be generated in order to train a classifier. Different methods have been used to generate negative peptides, *e.g*. through sequence shuffling (16), sampling from the human proteome (13) or through random sampling from uniprot (12, 15). Nevertheless, each method proposes certain assumptions about the nature of negative peptides and limits the kind of models that can be trained. For example, using sequence shuffling prevents the model from exploiting the cystine-depletion in MS-based inferred bound peptides, *i.e*. the positives, (31, 32). On the other side, omics integration and annotation with shuffled peptides is not trivial as these peptides are not obtained or derived from a real biological sequence. Further, sampling negatives from uniprot assumes that only peptide sequences are governing presentation and does not account for sequence-independent protein features and as a result renders multimodal training impractical. Finally, sampling from the human proteome assumes that the observed set of peptides are the full immunopeptidome and all non-observed peptides are non-presented. As shown above, the set of presented peptides is bounded by the sensitivity of the mass spectrometry and thus, the definition of negatives depends on the sensitivity of mass spectrometry. As a result, future effort shall focus on improving the sensitivity of the mass spectrometry to enable a more complete characterization of the immunopeptidome.

In conclusion, here we demonstrated an important role for sequence-independent features, *e.g*. the transcriptomic landscape of the cell and subcellular location, in shaping HLA-II immunopeptidomes. Further, modelling sequence dependent and independent features using deep multimodal models enables a more accurate prediction of *in vivo* presented peptides as illustrated with *PIA-M*. However, we have mainly focused on modelling and characterizing the rules governing the presentation of self-proteins which might differ from the rules governing the presentation of microbial peptides. Hence, future research shall focus on disentangling sequence-independent features governing microbial presentation by HLA-II proteins *in vivo*. Furthermore, the impact of peptide presentation on immunogenicity, immunodominance and T cell activations remains poorly understood. Hence, investigating the effect of different facets of peptide presentation, for example, tissue-restricted presentation and T cell activation, is a promising future direction to improve our understanding of the immune system.

## AVAILABILITY

The multi-omics linking toolkit (*OmLiT*) is an open-source library that is available under a permissive MIT-licence in the GitHub repository (https://github.com/ikmb/OmLiT). Meanwhile, the pipeline along with the pretrained models are available at (https://github.com/ikmb/PIA-inference). Finally, the web server for running predictions using the trained models is available at (https://hybridcomputing.ikmb.uni-kiel.de/pia).

## SUPPLEMENTARY DATA

Supplementary Data are available at NAR online

## ACKNOWLEDGEMENT

We are truly thankful to *Carina Saggau* (*Institute of Immunology; UKSH, Kiel, Germany*) and *Corinna Bang* (*Institute of Clinical Molecular Biology; Christian-Albrechts-Universität, Kiel, Germany*) for their support of this work.

