## Supplementary figures for "Predicting Peptide HLA-II Presentation Using Immunopeptidomics, Transcriptomics and Deep Multimodal Learning"

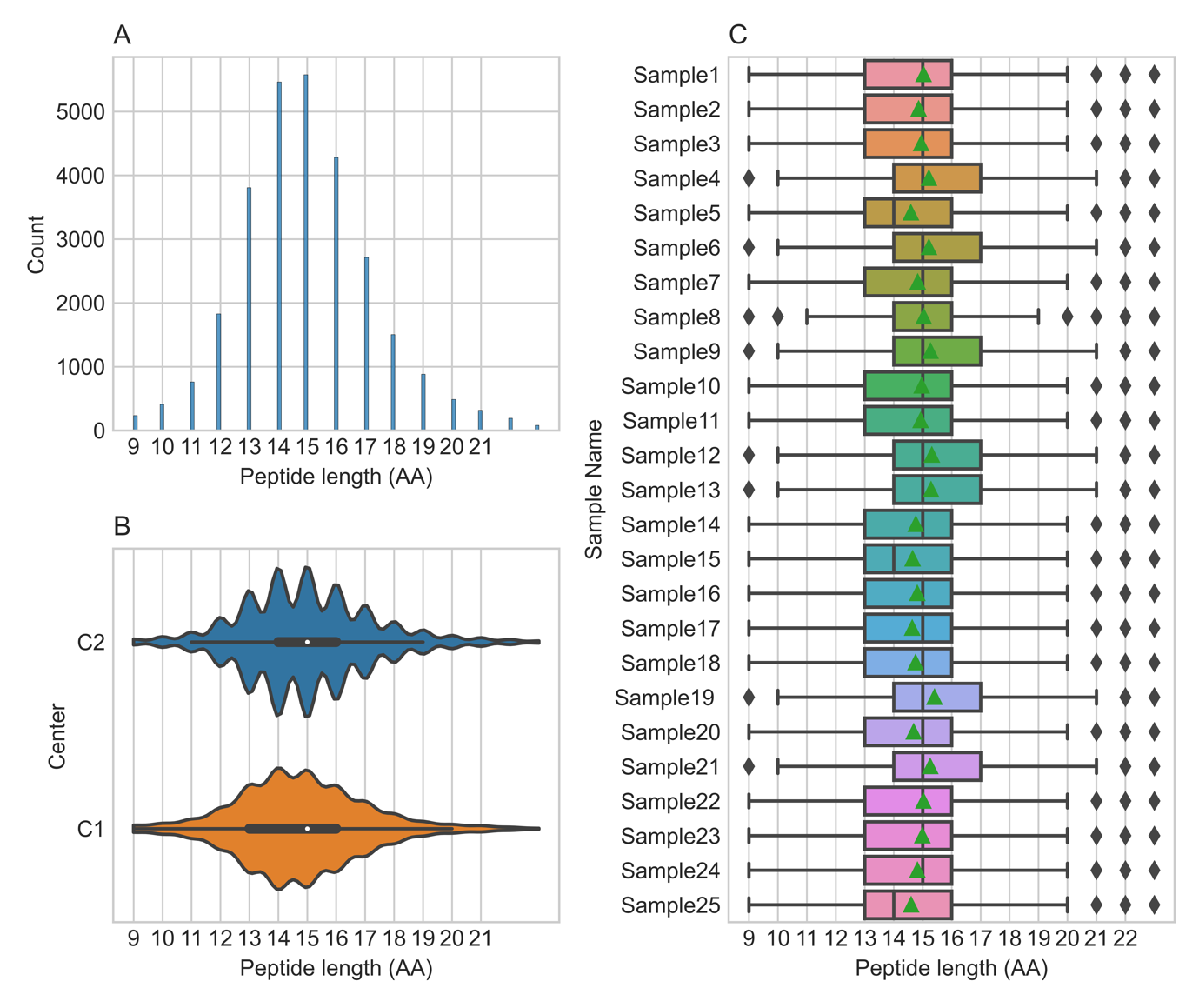

**Figure S1: Peptide length distribution for all peptides identified in the 25 samples measured in the current study.** (**A**) A histogram showing the count of peptides identified with a specific length, e.g. 13-mers or 15-mers using all datapoints. As shown in the figure, most peptides are 13-mers to 17-mers. (**B**) A comparison between the length distribution of all peptides measured in the two-proteomics centers namely, center 1 (C1) and center 2 (C2) as described in the Materials and Methods section. (**C**) Peptide length distribution per sample where black lines represent the median while green triangles represent the mean.

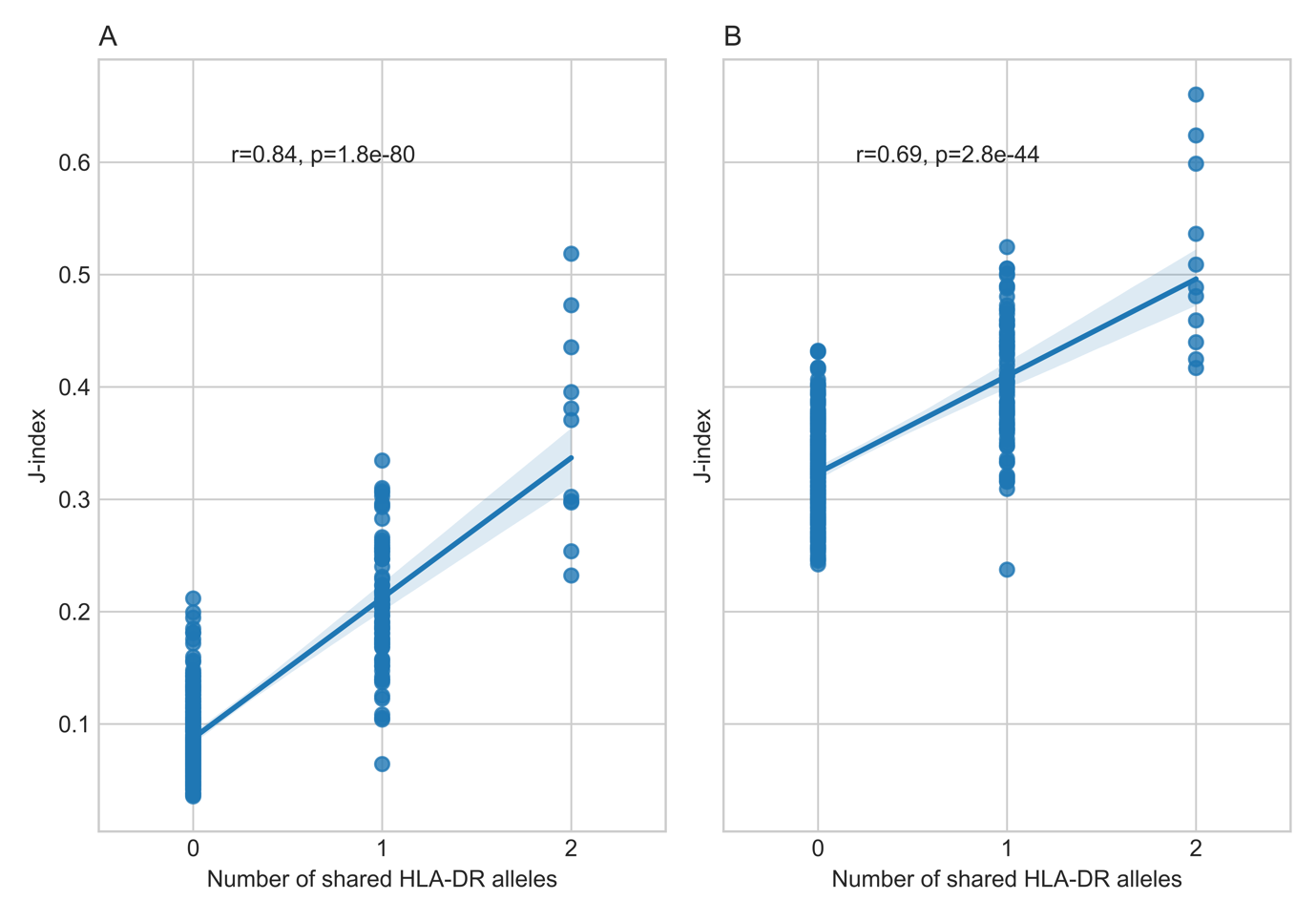

**Figure S2: The relationship between the number of shared HLA-DR alleles and J-index between all unique sample pairs, r is the person correlation coefficient while p is the P-value. (A**) shows the relationship between the number of shared HLA-DR alleles and J-index computed at the peptide-level while (**B**) shows the same relationship but at the protein-level.

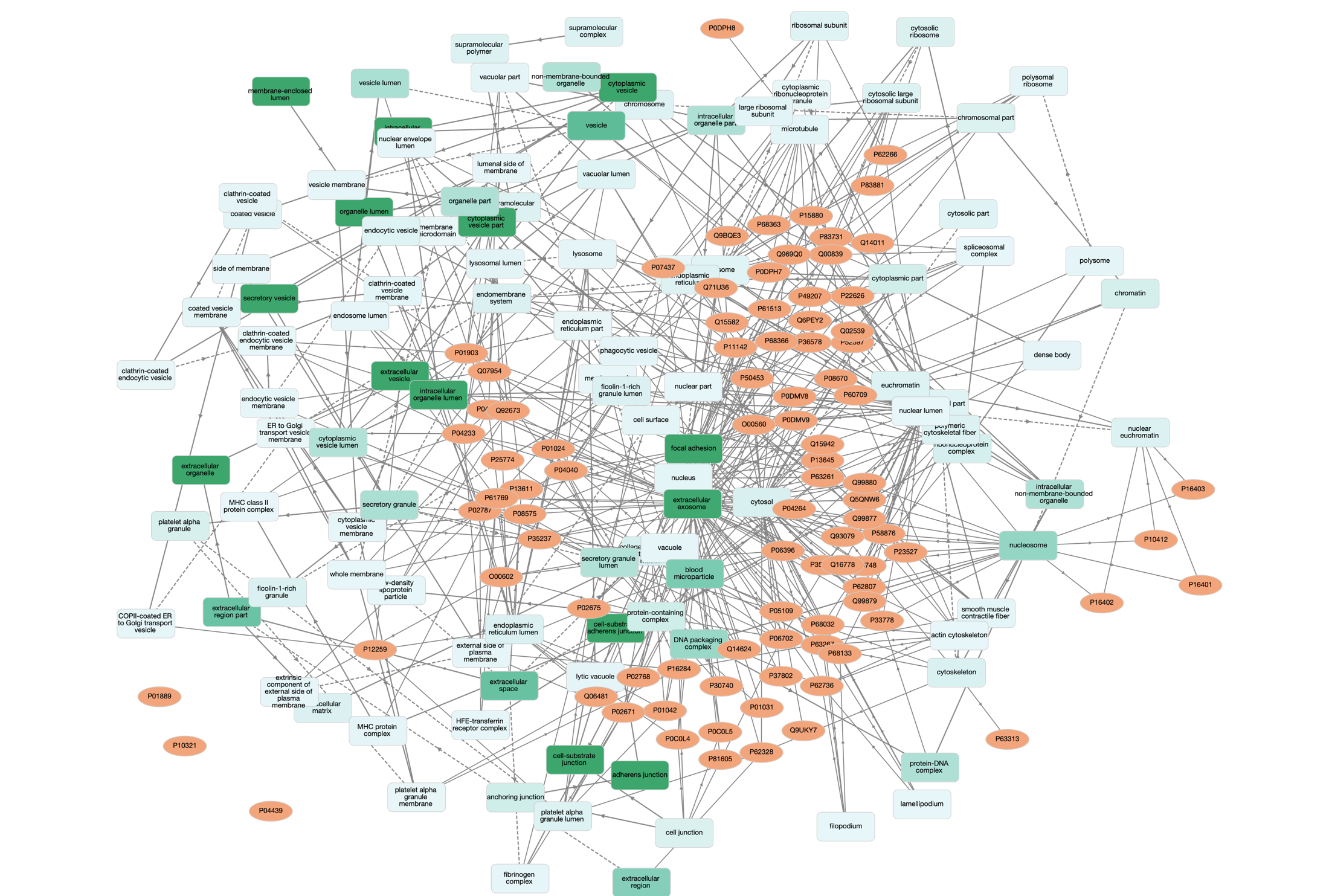

**Figure S3: Visualization of gene ontology enrichment analysis (GOEA) focusing on the proteins presented in all samples measured in the current study.** Gene network has been conducted using GOnet web-application (<https://tools.dice-database.org/GOnet/>) (1).

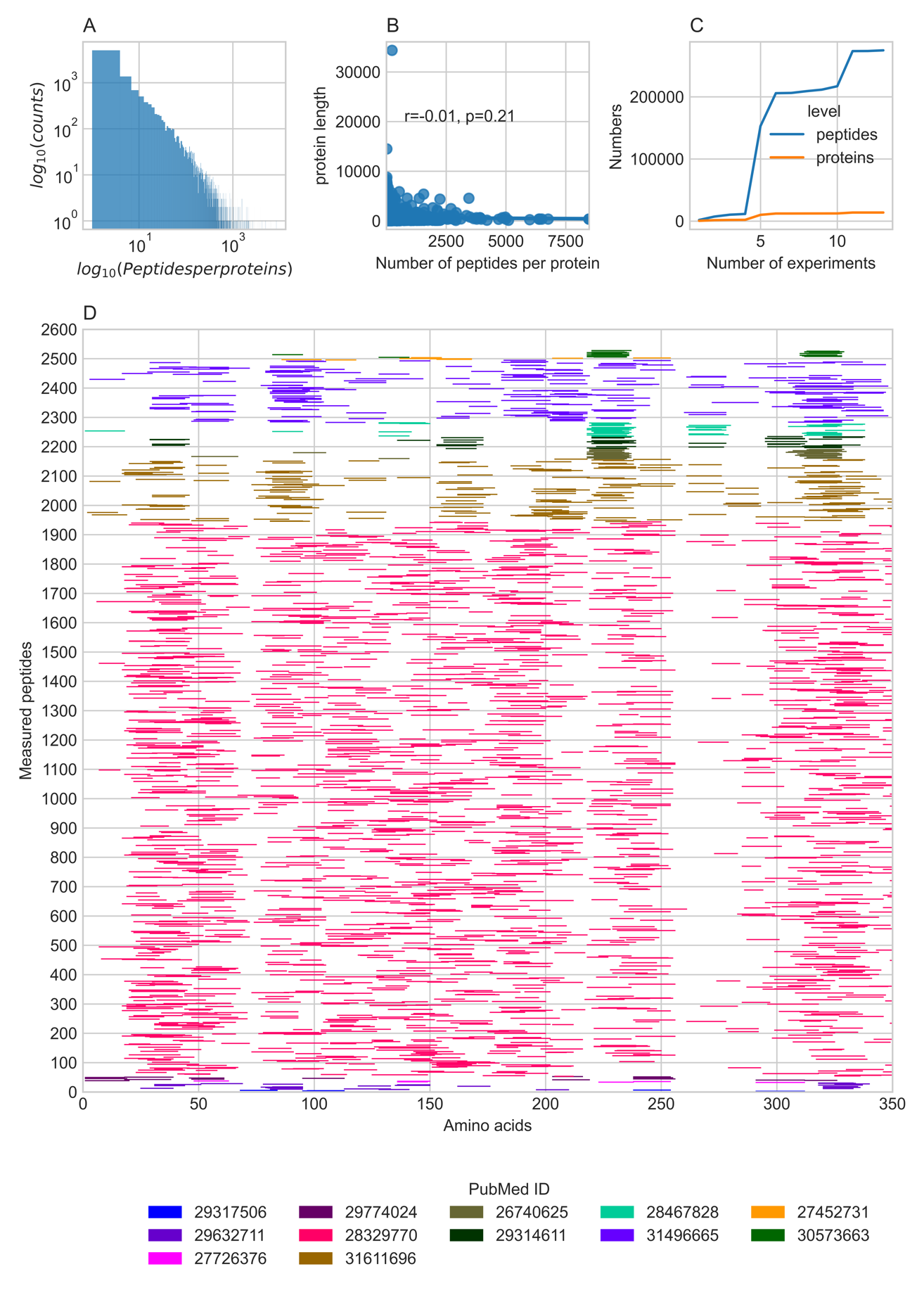

**Figure S4: A summary of publicly available immunopeptidomics datasets. (A) A histogram for the average number of peptides per proteins across all datasets**. (**B**) The correlation between the number of peptides per proteins and protein length across all datasets. (**C**) The cumulative number of unique peptides and proteins across different datasets. (**D**) A coverage representation for the protein with most peptides Actin, cytoplasmic 1 (ACTB) across all datasets. The X-axis represents the protein backbone while the y-axis represents the total number of unique peptides identified in different datasets. Peptides are aligned to the reference protein sequence and are visualized as short rectangles colored according to source publication; color keys represent the PubMed ID of each study.

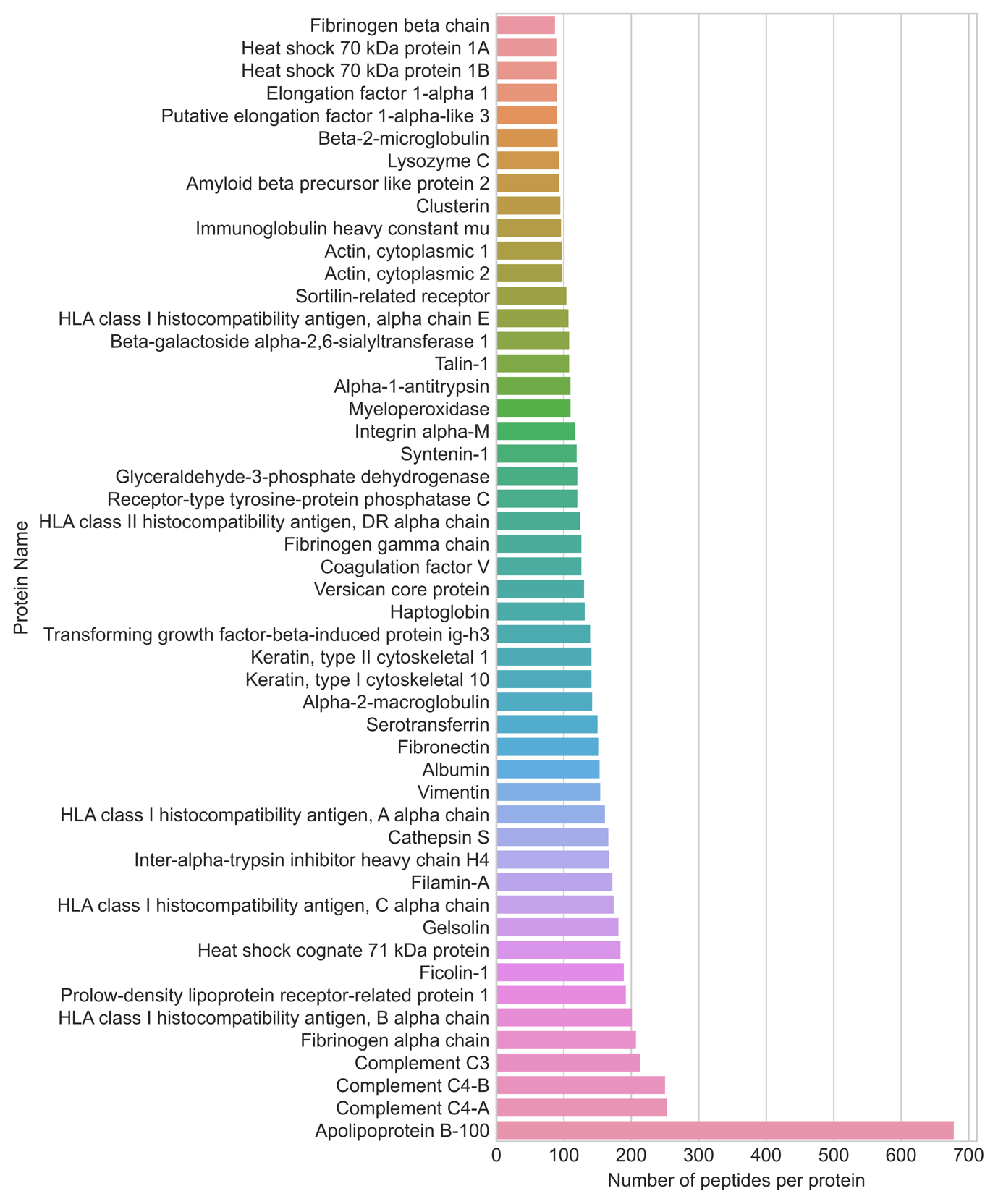

**Figure S5: Top-50 most presented human proteins in the dataset generated in the current study where presentation is measured as the number of peptides per protein.**

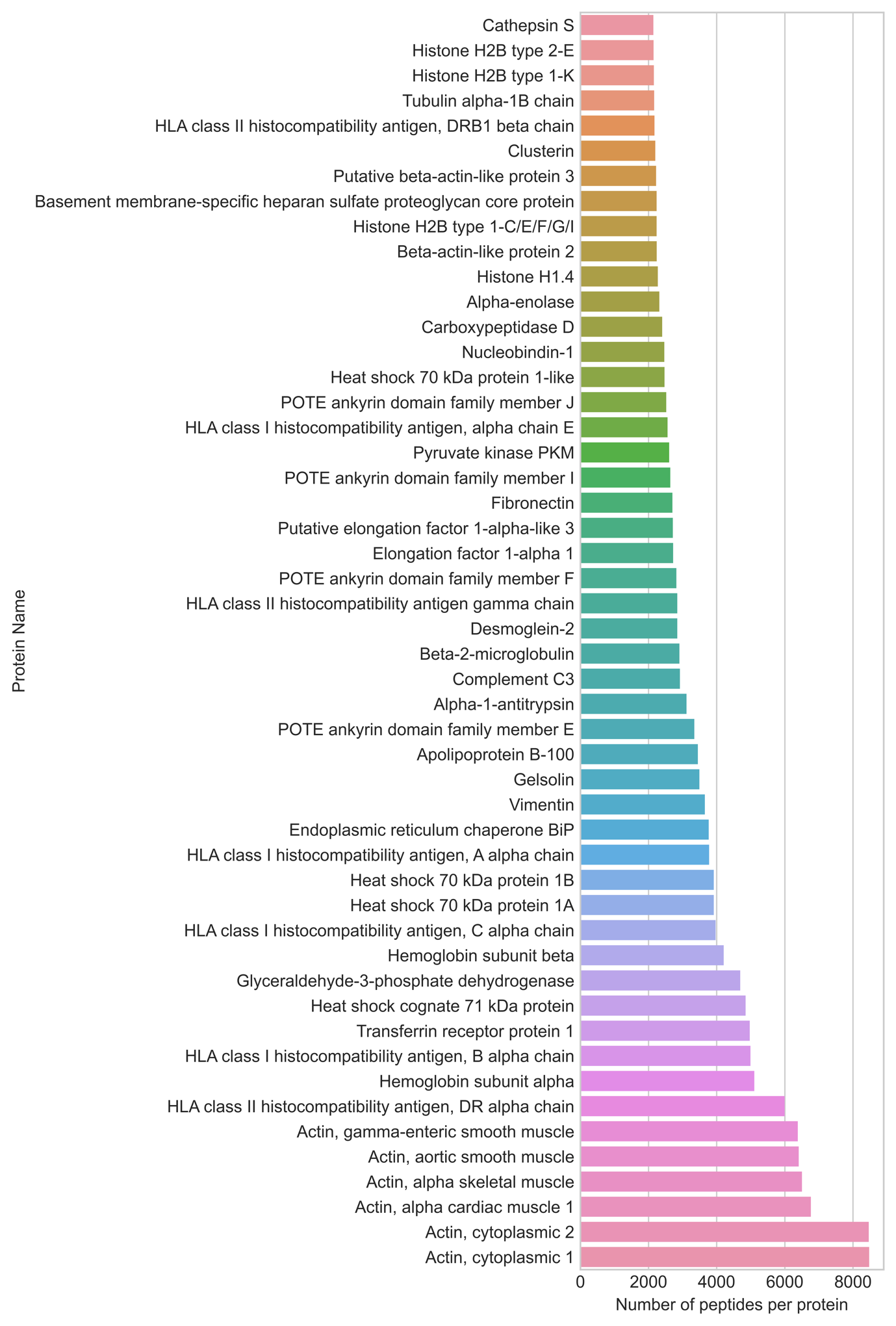

**Figure S6: Top-50 most presented human proteins in the public database where presentation is measured as the number of peptides per protein.**

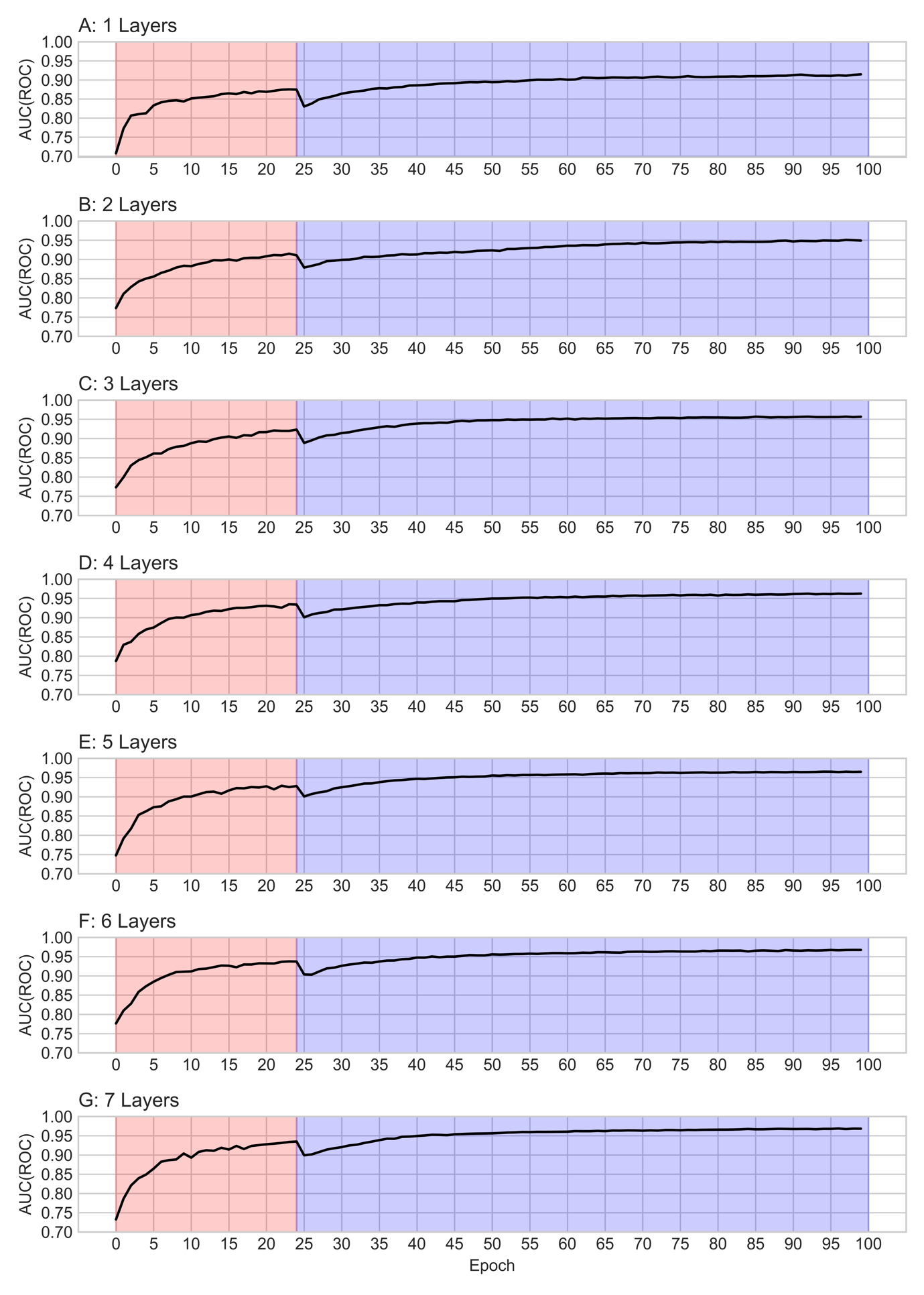
**Figure S7: The performance of different models with an increasing number of layers.** The x-axis illustrates the number of training epochs while the y-axis illustrates the area under the receiver operator curve (AUC(ROC)) calculated on the test dataset. The red area highlights training using the single allele dataset while the blue section highlights training the on the full dataset with allelic deconvolution.

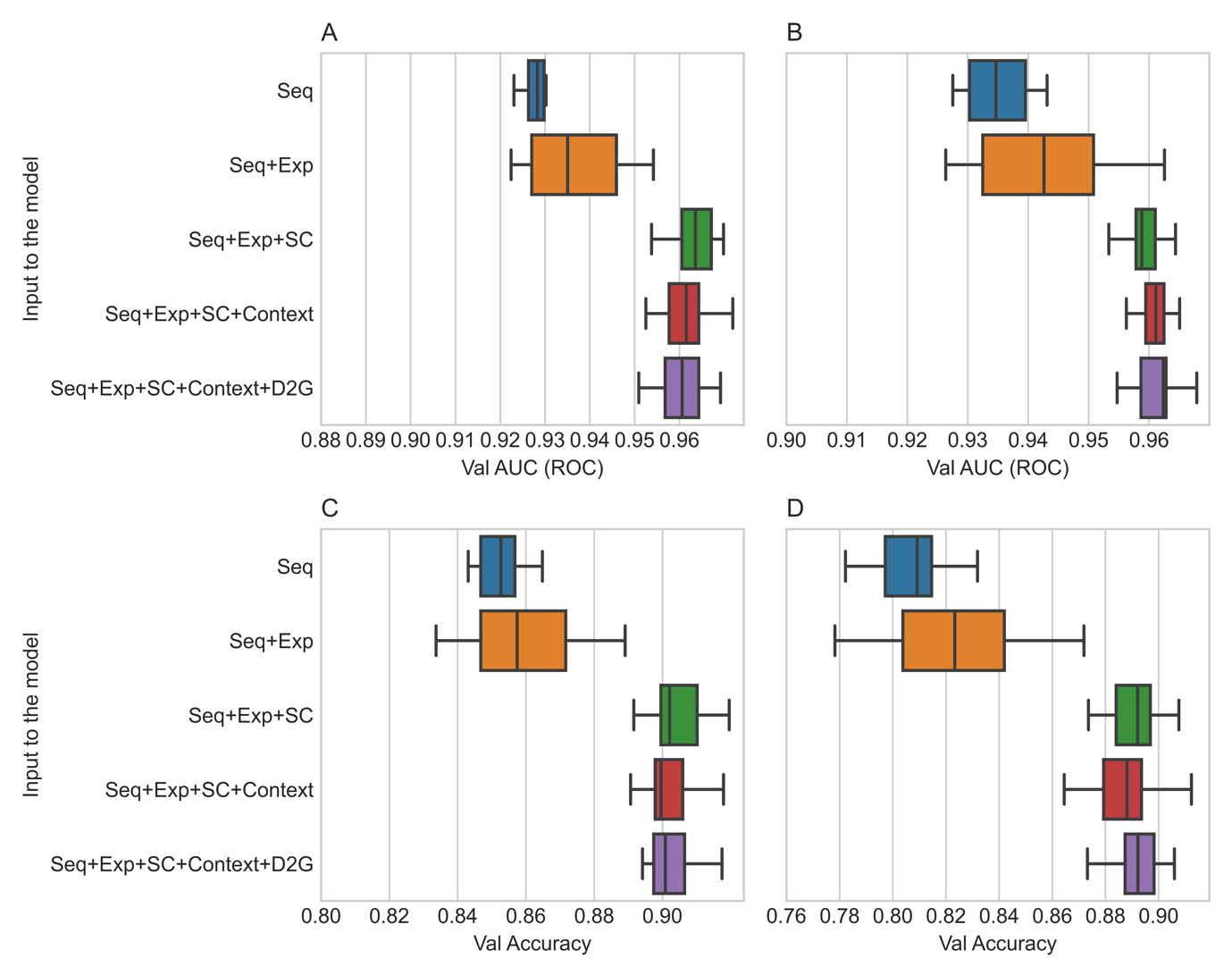

**Figure S8: The performance of multimodal training using two immunopeptidomics datasets generated in the current study, the first is from a homozygote HLA-DRB1*04:01 sample (Sample13) while the second is from an HLA-DRB1*15:01 sample (Sample6).** Seq stands for model trained on the peptide sequence only, Seq+Exp stands for models trained on the peptide sequence and parent expression level. While Seq+Exp+SC stands for models trained on the peptide sequence, parent expression level and encoded subcellular location, meanwhile Seq+Exp+SC+Context stands for models trained on the peptide sequence, parent expression level, encoded subcellular location and a context vector representing the expression level of all protein-coding genes in the tissue. Finally, Seq+Exp+SC+Context+D2G is a model trained on peptide sequence, parent expression level, encoded subcellular location, context vectors along with the distance to the nearest glycosylation site. (**A**) A comparison between the performance of different multimodal models on the validation dataset of Sample6 using the area under the receiver operator curve (AUC (ROC)) as an evaluation metric. (**B**) A comparison between the performance of different multimodal models on the validation dataset of Sample13 using the area under the receiver operator curve (AUC (ROC)) as an evaluation metric. (**C**) A comparison between the performance of different multimodal models on the validation dataset of Sample6 dataset using binary accuracy as an evaluation metric. (**D**) A comparison between the performance of different multimodal models on the validation dataset of Sample13 using binary accuracy as an evaluation metric.

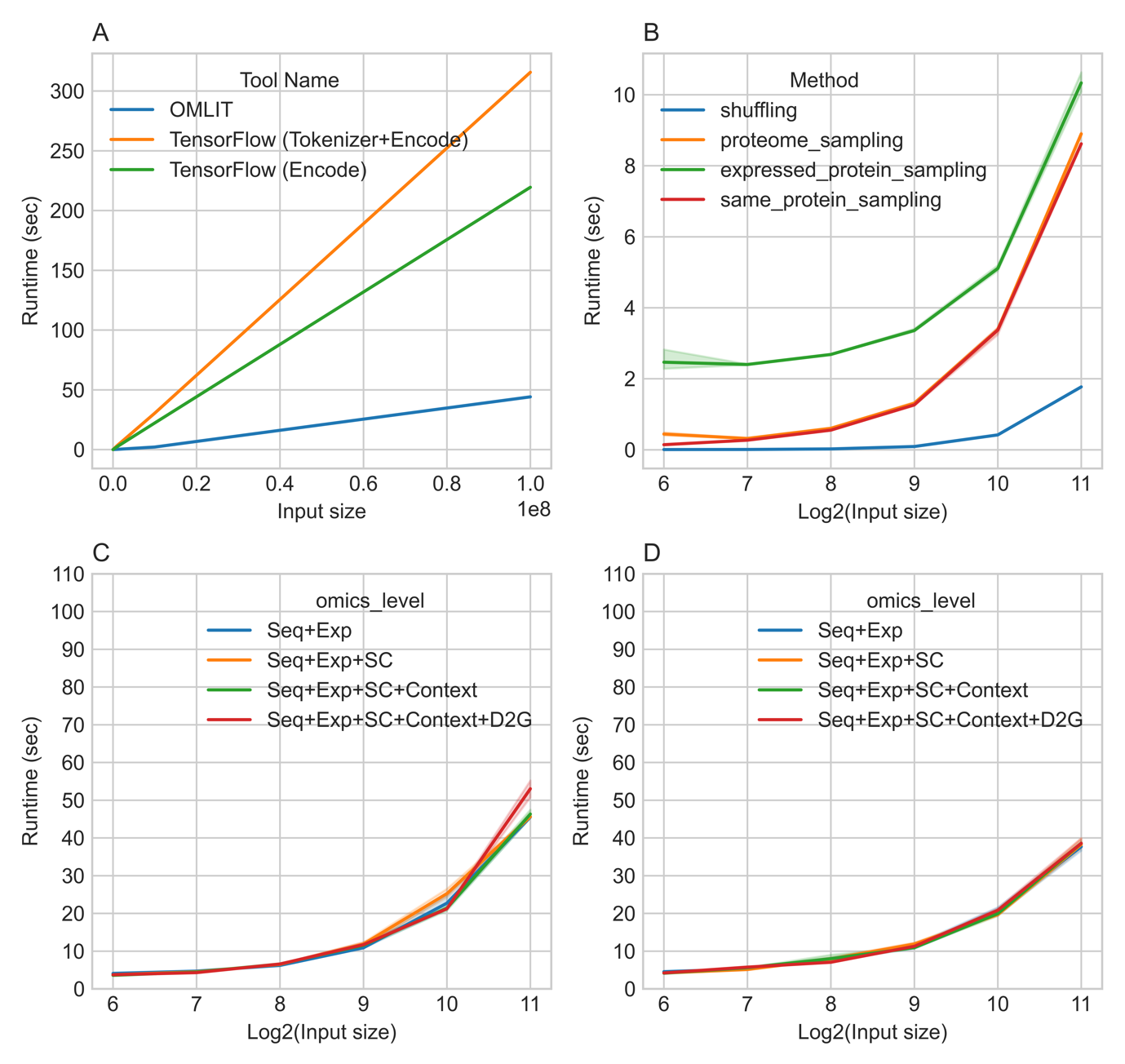

**Fig. S9: The performance of the omics linking toolkit (OmLiT) library.** (**A**) a comparison between the encoding speed of OmLiT relative to the Keras tokenizer of TensorFlow with an increasing number of samples. (**B**) The run time of different negative generation algorisms implemented in OmLiT where shuffling stands for shuffling the sequence of positive amino acids, while proteome sampling stands random sampling from the human proteome, expressed protein sampling stands for random sampling from the expressed proteins in a specific tissue and finally, same protein sampling stands for sampling from the same protein where the positive peptide was observed. (**C**) The OmLiT runtime while annotating different omics layer, Seq stands for model trained on the peptide sequence only, Seq+EXP stands for models trained on the peptide sequence and parent expression level. While Seq+Exp+SC stands for models trained on the peptide sequence, parent expression level and encoded subcellular location, meanwhile Seq+Exp+SC+Context stands for models trained on the peptide sequence, parent expression level, encoded subcellular location and a context vector representing the expression level of all protein-coding genes in the tissue. Finally, Seq+Exp+SC+Context+D2G is a model trained on peptide sequence, parent expression level, encoded subcellular location, context vectors along with the distance to the nearest glycosylation site. In (**C**) only one allele was used in the benchmarking study while in (**D**) the benchmarking dataset was simulated to contain five alleles. OmLiT splits the input data by allele and annotate data from each allele on parallel using a pool of worker threads. Hence, execution time was lower in D relative to C given that both had the same input size.

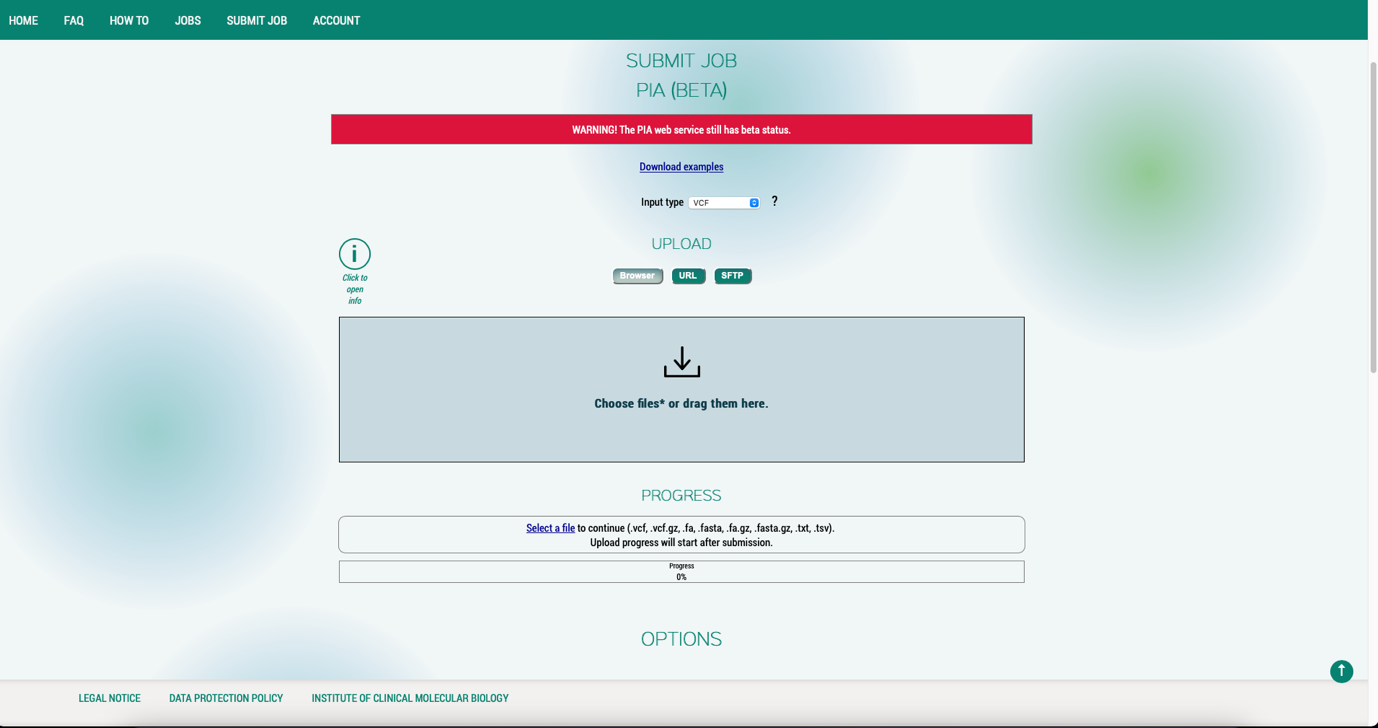

**Figure S10: An overview of the developed web interface. Here, the user has first to choose the input type which can be an input table, a genetic table, an VCF file or a FASTA file along with uploading these inputs to the webpage.** Subsequently these files are intercepted and subsequently fed to the PIA-P pipeline. The GitHub page of the project provides a detailed description of the required formatting of the inputs along with an in depth description of the execution logic with each input type.
