## Supplementary Methods for "Predicting Peptide HLA-II Presentation Using Immunopeptidomics, Transcriptomics and Deep Multimodal Learning"

**PBMC and sample isolation**

Blood samples were obtained from the University Medical Center Schleswig-Holstein (UKSH) blood bank as leukocyte reduction system (LRS) cones. Peripheral blood monocytes (PBMCs) were isolated from the cones using Ficoll-Hypaque centrifugation. Cells were counted using propidium iodide staining on a MACSQuant analyzer (Miltenyi Biotec). Cell pellets were snap-frozen using liquid nitrogen and stored at -80 °C.

**HLA-DR pulldown**

Briefly, the PBMCs were incubated with 2 ml of lysis buffer containing 0.5% IGEPAL CA-630, 0.25% Sodium deoxycholate, 1 mM EDTA in PBS, pH 7.5 (at 4°C), 1mM phenylmethylsulfonyl fluoride (PMSF), 0.2 mM iodoacetamide, and complete protease inhibitor cocktail (*Ultra Roche*) for 1 hour. Next, Poly-Prep columns (*BioRad*) were prepared for HLA pull down, by incubation with 200 μL of L243 antibody (*BioXCell*) coupled with AminoLink Plus resin (*ThermoFisher*) and washing with 10 ml lysis buffer. Subsequently, 7 ml of lysis buffer were added to the column followed by the addition of 2 ml of lysed cells. Next, the column was sealed with a paraffin film and the content of the column were incubated on a rotator for 2 hours at 4°C. Then the resin was first washed with 10 ml of buffer containing 150 mM sodium chloride and 20 mM Tris, followed by 10 ml of buffer containing 400 mM sodium chloride and 20 mM Tris, after that it was washed with a 10 ml buffer containing 150 mM sodium chloride and 20 mM Tris and lastly with a 10 ml buffer containing 20 mM Tris in LC-MS *LiChrosolv* water. Subsequently, peptides were eluted using 10% acetic acid. The eluted peptides were then washed with 50% acetonitrile and 0.1% TFA, and subsequently purified using C18 columns. Finally, the washed peptides were dried using lyophilization and then stored at -20°C.

**Mass spectrometry measurement and peptide identification**

Mass spectrometry measurements for C1 and C2 were conducted using the same settings described previously by *ElAbd* *et al.* (1)*.* Peptides were identified using the *MHCquant* (2) pipeline revision 1.5.1. The human reference proteome obtained from *UniProt* (3) (accession number: UP000005640) was used for peptide identification. The file was filtered to contain only proteins defined or included in the *SWISS-PROT* (4) database. After filtration, peptide mapping was conducted against 20,360 proteins. Further, peptide length was constrained to be between 9 and 23 amino acids and the false discovery rate (FDR) was set to 1%. Finally, data analysis and integration was conducted using *IPTK* (1) version 0.6.21.

**RNA-sequencing and transcriptomic analysis**

RNA extraction and purification along with sequencing was conducted at the Competence Centre for Genomic Analysis (CCGA) Kiel using two replicates per sample as written above. Sequencing was conducted using 150 bps paired-end sequencing and on average ~100 million reads were produced per sample. Read alignment and expression quantification was conducted using the *nf-core*/*rnaseq* pipeline (version 3.2) (5, 6). Public gene expression data was obtained from the Human Protein Atlas ([https://www.proteinatlas.org](https://www.proteinatlas.org/)) (7).

**HLA genotyping**

In total 23 individuals were genotyped on Illumina’s Global Screening Array (GSA) version 3.0. Data were checked for general quality metrics such as genotype call rate per sample and SNP as well as sample ancestry. The average SNP genotype call rate for all individuals in the HLA region (chromosome 6: 29-34Mb) was 99.1% and the median genotype call rate per individual was 0.06%. We extracted pre-imputation SNP genotypes from the HLA region and used them as input for HLA genotype prediction with the random-forest-based machine learning tool HIBAG (version 1.20.0)(8). Imputation of the classical histocompatibility leukocyte antigen (HLA) class II locus HLA-*DRB1* was performed using the multi-ethnic reference model published by Degenhardt *et al.* (9) which was modified to use the variants available on the GSA. Marginal posterior probabilities for single HLA alleles across all predicted HLA genotypes were computed as described in Degenhardt *et al.* (10). In the end, only alleles of the *DRB1* locus were used for analysis.

**Feature encoding**

Next to the peptide sequence and the HLA allele we included the RNA expression level, the distance to the nearest glycosylation site and the subcellular compartments as model features along with the expression value of all protein coding genes in the cell. RNA expression data were either experimentally measured as discussed above or obtained from the Human Protein Atlas (7). During model training and HLA-peptide-binding inference, RNA expression levels (measured in transcripts per million (TPMs)) were log-transformed as ${log}_{10}\left( expression level+1 \right)$ to stabilize training and accelerate convergence. To calculate the distance to the nearest glycosylation site for a peptide in its parent protein, the parent protein annotations were downloaded from *Uniprot* (19) using *IPTK* (1) and the absolute distances between the starting site of the peptide and all glycosylation sites in the protein were calculated. After that, the minimum distance was reported as the nearest glycosylation site. In case the parent protein did not have any reported glycosylation sites, its full length was considered as the nearest glycosylation site. Meanwhile, if within the peptide boundaries in the parent protein a particular glycosylation site was reported, then the nearest glycosylation site was set to zero. Lastly, the calculated distance was log-transformed as ${log}_{10}\left( distance to the nearest glycosylation site+1 \right)$ as discussed above.

The subcellular compartments of each protein were encoded in a ‘multi-hot’ fashion using data obtained from the gene ontology consortium (11, 12). Briefly, a vector with the same length as the number of unique compartments defined by the consortium was first allocated and filled with zeros. Next, each cellular compartment was assigned an index in the vector, then, for each protein cellular compartments were extracted and their corresponding indices in the allocated vector was assigned a value of one while all other indices were assigned a value of zero. Thus, enabling the generation of a fixed-length numerical representation of subcellular compartments. Lastly, amino acid encoding was conducted using an end-to-end embedding as discussed by *ElAbd et al*. (13) with the dimensionally of the embedding space set to 32.

**Modelling sequence-independent features**

Different sequence-independent features were modelled along the peptide sequence to quantify the effect of modelling these features on improving the predictive performance of the model. To this end, the immunopeptidome of two homozygote samples that were generated in the current study were used as development datasets for quantifying the impact of multimodal training on the performance. The first sample was obtained from a homozygous individual with an HLA-*DRB1*04:01* while the second was obtained from an HLA-*DRB1*15:01* homozygous sample. For both samples negative peptides were generated by sampling decoys from the set of human proteins that were not identified in the immunopeptidome of each sample and are also derived from genes with an expression level of at least 1 TPM in the PBMCs. Further, we annotated both negative and positive peptides with four sequence-independent features, namely, the expression level of the parent transcript (exp), the subcellular location of parent proteins (SC), the distance to the nearest glycosylation site (D2G). Finally, the expression level of all protein coding genes in the target tissue (Context) as defined in the Human Protein Atlas (7) (**RNA-sequencing and transcriptomic analysis**). Next, we trained different multimodal models on different combination of these features along with sequence-dependent features, *i.e.* peptide-sequence and HLA-II pseudo-sequence (14, 15), using 80% of the data and evaluated the performance on a 20% validation dataset.

**Development of omics linking toolkit (OmLiT)**

The omics linking toolkit (*OmLiT*) library was developed to accelerate the annotation and the encoding of multi-omics data for an input list of peptides. Briefly, the library is divided into two parts, a Rust execution engine, and a Python interface. All the encoding and the computationally heavy annotation tasks are conducted using the Rust execution engine where multithreading is used to accelerate the performance. The Python interface provides a thin wrapper around the execution engine and enables an easy integration with the rest of the training and inference infrastructure. Here, *Rayon* was used for implementing parallelism in the Rust-sided part, while *PyO3* was used to provide a Python-binder to the Rust-sided code, finally, the *NumPy* C API was used for moving the encoded data from the Rust-sided code to Python.

**Assembly of training datasets and PIA-S training**

***Assembly of datasets***

*PIA-S* was trained either using a large corpora of publicly available immunopeptidomic datasets (**Table S3**) following the same approach used by *Reynisson* and colleagues (16) with a total number of positive peptides equals to 715,642. Second, using this dataset combined with datasets generated in the current study (22 datasets from 20 donors (15 measured in C1 only, 3 measured in C2 only and 2 in C1 and C2 (4 datasets in total) were used for training) with a total number of positive peptides equals to 798,242. The remaining 3 datasets were used for benchmarking (**Materials and Methods**). For both training datasets (i) and (ii) negatives (*i.e.* peptides not observed to bind in the experimental setting) were sampled from the *SWISS-PROT* database (4) (available at *UniProt* (3)). Here, protein sequences shorter than 30 amino acids were removed. Then, for each positive peptide, five negative peptides with the same length as the positive peptide were randomly sampled from all proteins in the (4) database.

***Training***

After assembling the training dataset, it was randomly split into 80% training and 20% validation. Training was conducted in two stages: first using single-allele (SA) immunopeptidomics datasets, *i.e.* where peptides were eluted from a particular HLA protein, and second using the combination of SA datasets and multiple-alleles (MA) datasets. Where MA contains peptides eluted from multiple different HLA-II proteins, for example, different HLA-DR and HLA-DP proteins or two different HLA-DR proteins expressed at the surface of a heterozygous cell. If not stated otherwise, training on single HLA protein datasets was conducted for 25 epochs while training on the full dataset was conducted for 75 epochs. During the second phase of training, allelic deconvolution was implemented using a multiple-instance learning (MIL) framework recently described by *Cheng* and colleagues (17). In both phases, training was conducted in batches of size 4,096 examples with *Adam* (18) as an optimizer and with the binary cross-entropy as a loss function.

**Assembly of training datasets and PIA-M training**

***Assembly of datasets***

As explained above, *PIA-M* is a multi-modal architecture that utilizes multiple omics layers to predict presentation by HLA-II proteins. Here, the publicly available immunopeptidomics database described by *Reynisson* *et al.* (16) was used for training the models. Briefly, the identified peptide-sequences (database-wide) were mapped back against the human proteins identified at the *SWISS-PROT* (4) database to identify the parent protein(s) of each peptide using exact sequence matching. Next, the identified *UniProt* IDs were mapped back to ensemble transcript IDs using *IPTK* (1) which utilizes *UniProt* (3) REST-API for conducting the mapping step. After that, proteins that were mapped to an ensemble transcript without a measured expression level in the Human Protein Atlas (7) where removed (with all their associated peptides) from the database. Next, the distance to the nearest glycosylation site, the expression level of the parent transcripts in the target tissue and the subcellular locations of each parent protein were extracted from a precalculated database. This database was built to allow fast annotation of input peptides and proteins described above (***Feature encoding*).** Lastly, negatives were sampled from the human proteome using genes that had at least an expression level of 1 transcript per million (TPM) and were not defined in the set of presented proteins. Given that most studies included in the public database were obtained from different cell lines with a poor transcriptomic profiling in most cases, the expression profile of PBMC was taken as an approximation. Here, the ratio of positive to negative examples was one-to-five, meaning that for each positive peptide five negative peptides were also sampled.

***Training***

After assembling the training dataset, it was randomly split into 80% later used for training and 20% used for validation. Finally, training was conducted using batches each containing 4,096 examples with *Adam* as an optimizer for, if not stated otherwise, 100 epochs where the first 25 epochs were conducted using the mono-allelic data (as described above) while the last 75 epochs were conducted using the full dataset with allelic deconvolution being implemented following the algorithms described by *Cheng et al.* (17).

**Training protocol**

All neural networks were implemented using *Keras* (19) and *TensorFlow* version 2.7.0 (20). Training was conducted using a GPU node with eight Tesla V100-SXM2 GPUs each equipped with 32 GB of memory. The node also had 512 GB of RAM and an Intel® Xeon® Gold 6234 CPU. *Adam* (18) was used as an optimizer using the default learning rate and parameters as implemented in *Keras* version 2.7.0.

**Benchmarking the execution speed of PIA-S and PIA-M**

To measure the execution speed of *PIA-S* and *PIA-M* against other publicly available tools, the multi-modal (multi-omics) dataset used for training *PIA-M* was used. Briefly, an increasing number of peptides was randomly sampled from the list of training peptides and each tool was called for three times using the same input and the execution time was recorded. Given that *NetMHCIIpan 4.0* and *NetMHCIIpan 4.1* just differ in the amount of the training data, benchmarking the execution speed only focused on *NetMHCIIpan 4.0.* The benchmarking study was conducted using the GPU node used for training the models (**Model Training**).
